# The mitochondrion of *Plasmodium falciparum* generates essential acetyl-CoA for protein acetylation

**DOI:** 10.1101/2022.03.02.482627

**Authors:** Sethu C. Nair, Justin T. Munro, Alexis Mann, Manuel Llinás, Sean T. Prigge

## Abstract

Coenzyme A (CoA) biosynthesis is an excellent target for antimalarial intervention. While most studies have focused on the use of CoA to produce acetyl-CoA in the apicoplast and the cytosol of malaria parasites, mitochondrial acetyl-CoA production is less well understood. In the current study, we performed metabolite labeling experiments to measure endogenous metabolites in *Plasmodium falciparum* lines with genetic deletions affecting mitochondrial dehydrogenase activity. Our results show that mitochondrial acetyl-CoA biosynthesis is essential for parasite growth and identify a catalytic redundancy between the two main ketoacid dehydrogenase enzymes, both of which are able to produce acetyl-CoA. The activity of these enzymes is dependent on the lipoate attachment enzyme LipL2, which is essential for parasite survival solely based on its role in supporting acetyl-CoA metabolism. We also find that acetyl-CoA produced in the mitochondrion is essential for the acetylation of histones and other proteins outside of the mitochondrion. Taken together, our results demonstrate that the mitochondrion is an essential *de novo* source of acetyl-CoA and is required for *P. falciparum* protein acetylation critical to parasite survival.

## Introduction

Malaria is a debilitating global infectious disease^1^. Efforts to control the disease have been largely successful in the last decade as evidenced by a declining number of cases and fatalities^2^. However, the lack of an effective vaccine^3^ and emerging drug-resistance against all available treatment regimens^4^ highlight the need for new antimalaria strategies. Coenzyme A (CoA) biosynthesis has been shown to be an excellent target for antimalarial intervention^5–8^, largely because CoA is required for the formation of the central metabolite acetyl-CoA. Acetyl-CoA occupies a critical position in many metabolic processes including fatty acid biosynthesis, amino acid metabolism, and TCA cycle metabolism^9^. It also plays a role in cell signaling, primarily through the acetylation of histones and other proteins, and acts as a molecular sensor that controls different cellular fates and metabolic decisions^10–12^. In the human malaria parasite *Plasmodium falciparum*, 1,146 acetylated proteins have been identified including glycolytic enzymes, histones, and secreted proteins that could have important roles throughout the parasite life cycle^13^. Histone acetylation has been shown to have an important role in controlling transcriptional regulation and cellular differentiation in these parasites^14, 15^. Drugs targeting histone deacetylases have been used to treat clinical cases of malaria^16–21^, highlighting the importance of protein acetylation and its potential as a therapeutic target.

Despite the significance of acetylation reactions in malaria parasites, the primary *de novo* source of acetyl-CoA is still unclear. There are three pathways that should be able to synthesize acetyl-CoA in different subcellular compartments of the parasite (**Figure 1A**). The most studied pathway uses an apicoplast-localized pyruvate dehydrogenase (aPDH) enzyme that catalyzes conversion of glucose-derived pyruvate to acetyl-CoA^22, 23^. Acetyl-CoA generated in the apicoplast is thought to be the carbon source for fatty acid biosynthesis in the organelle^24, 25^, however, fatty acid biosynthesis enzymes^26–28^ and aPDH proteins^23, 29, 30^ have been shown to be dispensable during the blood stages of malaria parasite development. Another important source of acetyl-CoA could be from an acetyl-CoA synthetase (ACS) enzyme located in the nucleus^6, 29, 31–33^ (**Figure 1A**). But the fact that ACS requires recycled or supplemented sources of acetate raises the question of whether this pathway can produce a sufficient amount of acetyl-CoA to support parasite growth. The third proposed biosynthetic source is through a mitochondrial enzyme complex called the branched chain ketoacid dehydrogenase (BCKDH).

**Figure 1.**
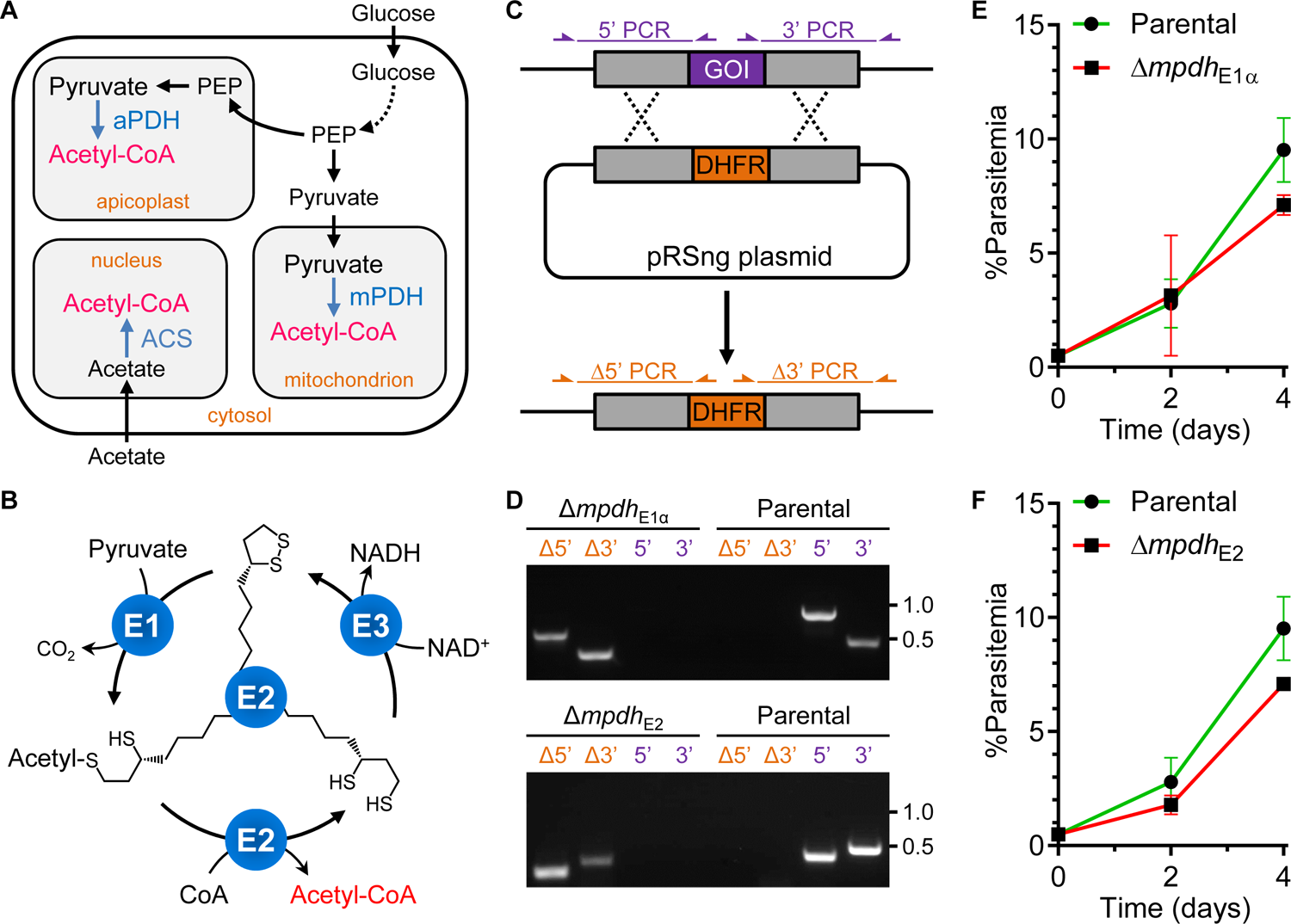
mPDH E1 and E2 subunits are not essential for blood stage parasite growth. **A.** Metabolic illustration showing possible acetyl-CoA biosynthetic pathways in the mitochondrion, apicoplast and nucleus of malaria parasites. PEP: phosphoenolpyruvate; aPDH: apicoplast pyruvate dehydrogenase; mPDH: mitochondrial pyruvate dehydrogenase; ACS: acetyl-CoA synthetase. **B.** Cartoon showing the conversion of pyruvate to acetyl-CoA by PDH. The enzymatic activities of the E1, E2 and E3 subunits of PDH are shown. Lipoic acid is covalently attached to the E2 subunit and is essential for PDH activity. **C.** Schematic representation showing how genes of interest (GOI) were replaced with a drug resistance cassette (DHFR) using the pRSng plasmid. The positions of genotyping PCR products designed to identify the wild type locus (purple) or the recombinant locus (orange) are shown. **D.** Genotyping PCR reactions confirming the deletion of *mpdh*_E1α_ (Top) and *mpdh*_E2_ (bottom). Based on the scheme shown in **C**, PCR amplicons demonstrate integration at the Δ5’ and Δ3’ loci and lack of parental parasites (as indicated by the failure to amplify at the wild type 5’ and 3’ loci). The parental line was used as a control. The primer sequences and amplicon lengths are described in **supplementary file 1**. **E, F.** Growth curves of Δ*mpdh*_E1α_ (**E**) and Δ*mpdh*_E2_ (**F**) parasites (red) compared to the parental NF54^attB^ line (green). Error bars represent the standard deviation from at least two independent experiments, each conducted in quadruplicate.

This enzyme complex is canonically thought to be involved in the catabolism of branched chain amino acids, but it may also be converting the pyruvate derived from glycolysis to acetyl-CoA (PDH activity)^34^. A study looking at the role of BCKDH in the murine malaria parasite (*Plasmodium berghei*) and a related apicomplexan parasite, *Toxoplasma gondii,* concluded that mitochondrial BCKDH is probably acting as a mitochondrial PDH (mPDH)^34^ and metabolic labeling studies using ^13^C-glucose in *P. falciparum* supported this notion by showing that the acetyl moiety of acetyl-CoA is derived from glucose^29^.

All known PDH enzymes require the enzyme cofactor lipoate^35^. Although malaria parasites have a *de novo* pathway to make lipoate in the apicoplast^36^, this pathway is dispensible^37–39^ and they cannot survive without scavenging lipoate^40^. Scavenged lipoate is exclusively used for lipoylating enzyme complexes in the mitochondrion, including BCKDH^40^, which has been localized to this organelle in *P. falciparum*^41^. Lipoylation of mitochondrial proteins appears to be essential since the expression of a bacterial lipoamidase enzyme that cleaves lipoate from mitochondrial enzyme complexes results in parasite death^42^. The toxic effects of lipoamidase were partially restored by acetate supplementation, suggesting that lipoylated enzyme complexes have a role in acetate metabolism^42^. Overall, several lines of evidence suggest that there is a pyruvate dehydrogenase in the parasite mitochondrion that uses lipoate as a cofactor and generates acetyl-CoA. However, the functional requirement of the scavenged lipoate and the role of lipoylated enzyme complexes in generating acetyl-CoA have not been fully elucidated.

In the current study, we investigated mitochondrial lipoylated enzyme complexes and their role in the biosynthesis of acetyl-CoA using a combination of genetic and metabolomic approaches. We demonstrated that BCKDH functions as a mitochondrial PDH in *P. falciparum* and we refer to it as mPDH throughout this manuscript. We also revealed functional redundancy between mPDH and α-ketoglutarate dehydrogenase (KDH), both of which are able to produce acetyl-CoA from glucose-derived pyruvate in the mitochondrion. Both dehydrogenase enzymes are dependent on lipoate as a cofactor and deletion of the lipoate attachment enzyme LipL2 completely blocks the production of acetyl-CoA from glucose. Parasites lacking LipL2 or lacking both mPDH and KDH can only survive when acetate is provided in the growth medium. We showed that the hypothetical ACS enzyme is responsible for the synthesis of acetyl-CoA from exogenous acetate, but required millimolar concentrations of acetate to bypass the loss of the mitochondrial enzymes. These results showed that under normal growth conditions, the mitochondrion is the major *de novo* source of acetyl-CoA in blood-stage malaria parasites. We also found that the mitochondrion acts as the source of acetyl-CoA used for the acetylation of histones and other proteins outside of the mitochondrion. In conclusion, this work explains the requirement for lipoate scavenging in blood-stage malaria parasites and reveals that the parasite relies on its mitochondrion for production of the essential metabolite acetyl-CoA.

## Results

### mPDH E1 and E2 subunits are not essential during the *P. falciparum* asexual blood stages

Pyruvate dehydrogenase enzymes classically catalyze the conversion of glucose-derived pyruvate to acetyl-CoA in the mitochondria of eukaryotic cells. However, in malaria parasites, the sole pyruvate dehydrogenase (aPDH) enzyme is not localized to the mitochondrion, but rather to a non-photosynthetic chloroplast organelle, the apicoplast, where acetyl-CoA is used to generate fatty acids (**Figure 1A**)^22, 23^. Gene deletion studies in the murine malaria parasite *P. berghei* and labeling studies in *P. falciparum* suggest that a mitochondrial Branched-chain Ketoacid Dehydrogenase (BCKDH) enzyme is most likely acting as a mitochondrial Pyruvate Dehydrogenase (mPDH)^29, 34^. mPDH enzymes are composed of multiple copies of three major subunits: E1, E2, and E3^35^ (**Figure 1B**). The E3 subunit (PF3D7_1232200) is not specific to mPDH and has been proposed to also function as part of the mitochondrial α-Ketoglutarate Dehydrogenase (KDH) and glycine cleavage system^43^. The E1 subunit of the mPDH complex is a heterodimeric protein (α [PF3D7_1312600] and β subunits [PF3D7_0504600]) that catalyzes the decarboxylation of pyruvate. The acetyl group released from the decarboxylation reaction is conjugated to a lipoate cofactor covalently attached to the E2 subunit (PF3D7_0303700) and is then transferred to CoA to form acetyl-CoA. To prepare for the next catalytic cycle, the E3 subunit oxidizes the lipoate cofactor, reforming the dithiolane ring^35^ (**Figure 1B**).

To address the role of mPDH during *P. falciparum* asexual development, we targeted the catalytic mPDH E1α subunit for deletion using Cas9-based genome editing (**Figure 1C, D**). Forward genetic screens predict that this protein is required for normal parasite growth (**Table 1**)^44, 45^. To our surprise, the resulting *Δmpdh_E1α_* line did not have a noticeable growth phenotype (**Figure 1E**) even though this protein was previously shown to be essential in *P. berghei*^34^. To validate this result, we generated a parasite line with a deletion in the gene encoding the E2 subunit, which should form the central core of the mPDH complex^35^. Again, consistent with the result observed for deletion of the E1α subunit, we did not observe a significant growth defect in the *Δmpdh_E2_* deletion line (**Figure 1F**). These results demonstrate that the mPDH enzyme complex is dispensable in blood stage *P. falciparum* parasites and does not play an essential role in acetyl-CoA biosynthesis or any other metabolic reaction.

**Table 1.**
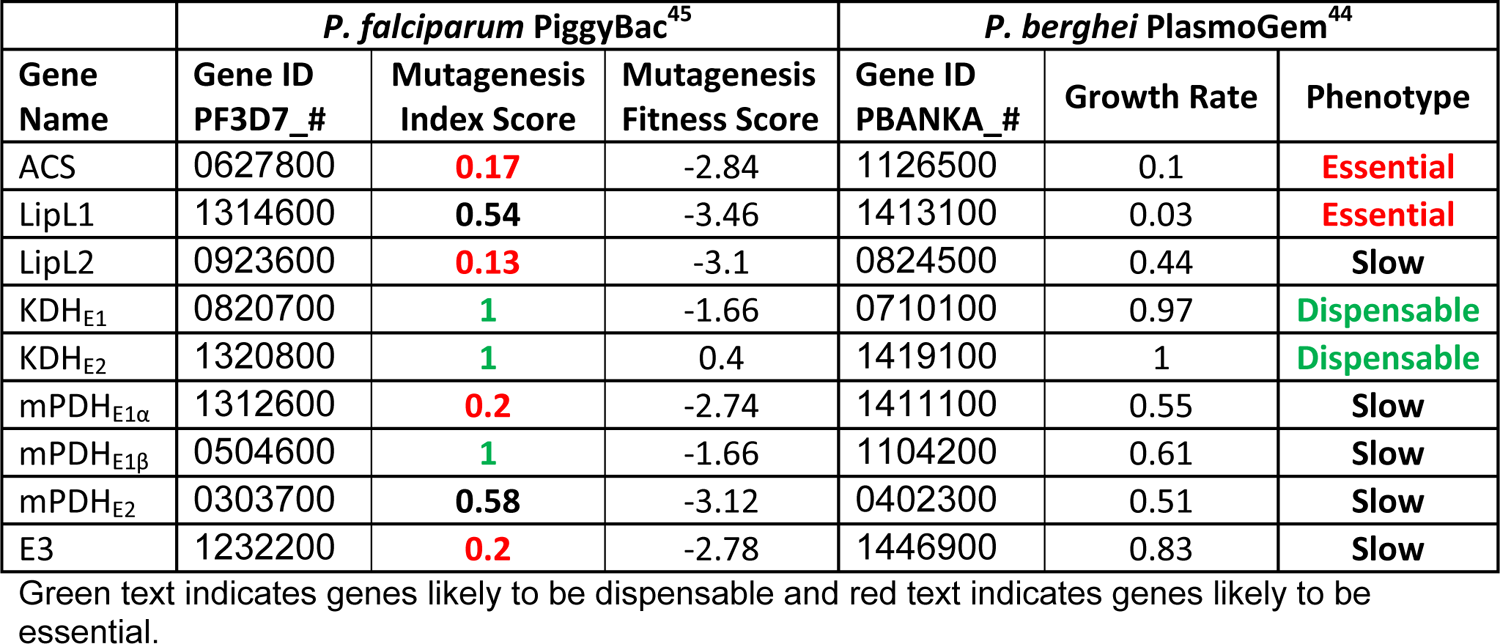
Predicted essentiality of genes based on forward genetic screens.

|  | *P. falciparum* PiggyBac<sup>45</sup> |  |  | *P. berghei* PlasmoGem<sup>44</sup> |  |  |
| --- | --- | --- | --- | --- | --- | --- |
| Gene Name | Gene ID PF3D7_# | Mutagenesis Index Score | Mutagenesis Fitness Score | Gene ID PBANKA_# | Growth Rate | Phenotype |
| ACS | 0627800 | 0.17 | -2.84 | 1126500 | 0.1 | Essential |
| LipL1 | 1314600 | 0.54 | -3.46 | 1413100 | 0.03 | Essential |
| LipL2 | 0923600 | 0.13 | -3.1 | 0824500 | 0.44 | Slow |
| KDH<sub>E1</sub> | 0820700 | 1 | -1.66 | 0710100 | 0.97 | Dispensable |
| KDH<sub>E2</sub> | 1320800 | 1 | 0.4 | 1419100 | 1 | Dispensable |
| mPDH<sub>E1α</sub> | 1312600 | 0.2 | -2.74 | 1411100 | 0.55 | Slow |
| mPDH<sub>E1β</sub> | 0504600 | 1 | -1.66 | 1104200 | 0.61 | Slow |
| mPDH<sub>E2</sub> | 0303700 | 0.58 | -3.12 | 0402300 | 0.51 | Slow |
| E3 | 1232200 | 0.2 | -2.78 | 1446900 | 0.83 | Slow |

Green text indicates genes likely to be dispensable and red text indicates genes likely to be essential.

### The mPDH E3 subunit is essential

As described above, the shared E3 subunit should also be required for KDH activity in the mitochondrion. However, previous deletion of the E1 subunit of KDH in *P. falciparum* strongly suggests that KDH activity is not essential during asexual blood-stage development^46^. Despite repeated attempts, we were unable to ablate the E3-encoding gene, suggesting that, even though the mPDH and KDH complexes individually are not required, the E3 subunit is essential. The E3 subunit may also be required for the activity of the glycine cleavage system H-protein, the only other lipoylated protein found in the parasite mitochondrion^47, 48^. Alternatively, mPDH and KDH may be capable of performing essential, but redundant functions.

Acetate bypass has been used previously in *P. berghei* to delete the mPDH_E1α_ subunit^34^ and bypass the toxic effects of a bacterial lipoamidase enzyme expressed in the parasite mitochondrion^42^. In both cases, an acetyl-CoA synthetase (ACS) enzyme (**Figure 2A**) was hypothesized to generate acetyl-CoA from acetate to enable parasite survival. Borrowing from this observation, we supplemented the culture medium with 5 mM acetate and were able to readily generate parasites lacking the E3 subunit (**Figure 2-figure supplement 1A**). Although the *Δe3* parasite line was reliant on acetate for growth (**Figure 2B**), growth arrest was not immediate, requiring several parasite growth cycles (**Figure 2-figure supplement 1B**). To further characterize this growth defect, we attempted to rescue parasite growth with acetate after various periods of deprivation. Acetate supplementation restored growth after 10 days of deprivation, but failed to restore growth after 12 days of acetate deprivation (**Figure 2C**). These results demonstrate that E3 is an essential protein and also suggest that E3 is ultimately required for acetate metabolism.

**Figure 2.**
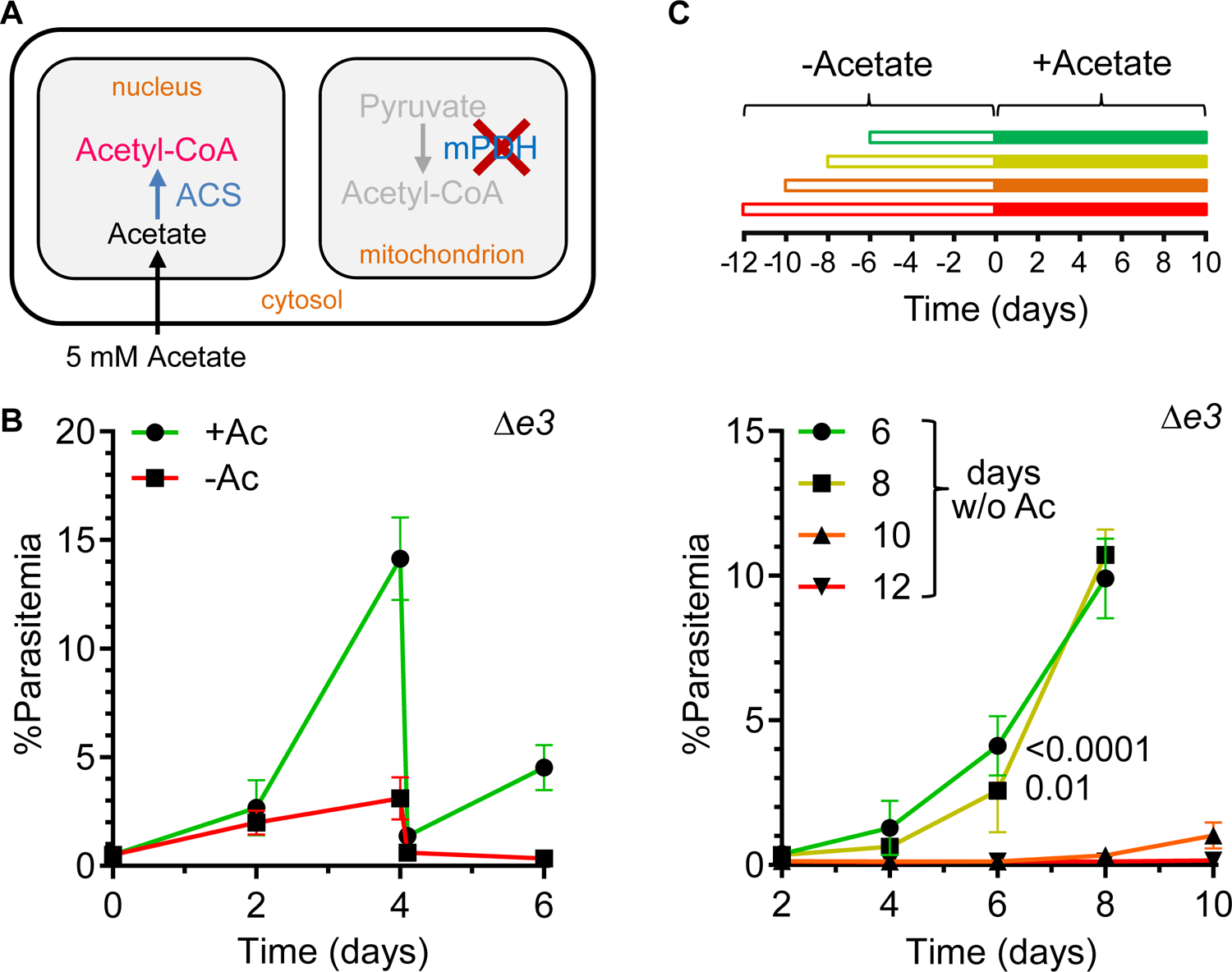
Δ*e3* parasites require acetate supplementation. **A.** Metabolic illustration showing how acetyl-CoA synthetase (ACS) could use exogenous acetate to generate acetyl-CoA, bypassing mitochondrial acetyl-CoA metabolism. **B.** Growth curves comparing the growth of Δ*e3* parasites in the presence and absence of 5 mM acetate (Ac). After four days, the cultures were diluted by a factor of 10 to avoid overgrowth. **C.** Survival of Δ*e3* parasites after different periods of acetate (Ac) deprivation. Parasite growth assays were initiated after different periods of maintenance in acetate-free medium (open bars in the schematic) followed by growth with acetate (solid bars). The plots below the schematic show the growth of these cultures after reintroduction of acetate. Significant parasite growth was observed on day six of the outgrowth period in the cultures deprived for 6 days or 8 days relative to the culture deprived for 12 days (two-way ANOVA, followed by Bonferroni’s correction; P values are provided in the plot). Error bars in the growth curves represent the standard deviation from at least two independent experiments, each conducted in quadruplicate.

### Mitochondrial ketoacid dehydrogenase activity is required for parasite survival

Biochemical studies have shown that KDH_E1_ is able to use pyruvate as a substrate, making it possible that KDH could function as a pyruvate dehydrogenase^49^. Although the KDH E1 subunit (PF3D7_0820700) is dispensable in blood stage *P. falciparum*^46^, it is possible that the KDH complex co-opts the E1 from mPDH since the complexes have been shown to swap subunits in other species^50^. Therefore, we tested if the KDH E2 subunit (PF3D7_1320800) is essential. Similar to the E1 knockout, *Δkdh_E2_* parasites did not display a significant growth phenotype compared to parental parasites (**Figure 3-figure supplement 1**). Taken together, the results show that the E1 and E2 subunits of KDH and mPDH are dispensable, but the shared E3 subunit is essential. This suggests that KDH and mPDH are functionally redundant. To explore this possibility, we used acetate supplementation and deleted the mPDH_E2_ subunit in the *Δkdh_E2_* parasite line, generating the *Δp_E2_/Δk_E2_* double deletion line (**Figure 3-figure supplement 2**). As anticipated, the *Δp_E2_/Δk_E2_* line required acetate supplementation for survival (**Figure 3A**), consistent with the hypothesis that KDH and mPDH are redundant. To characterize this phenomenon, we determined the acetate dependence of the *Δp_E2_/Δk_E2_* line and found that parasite growth is significantly compromised with 1.25 mM acetate or less (**Figure 3B**) and that the *Δp_E2_/Δk_E2_* parasites could survive twelve hours of acetate deprivation.

**Figure 3.**
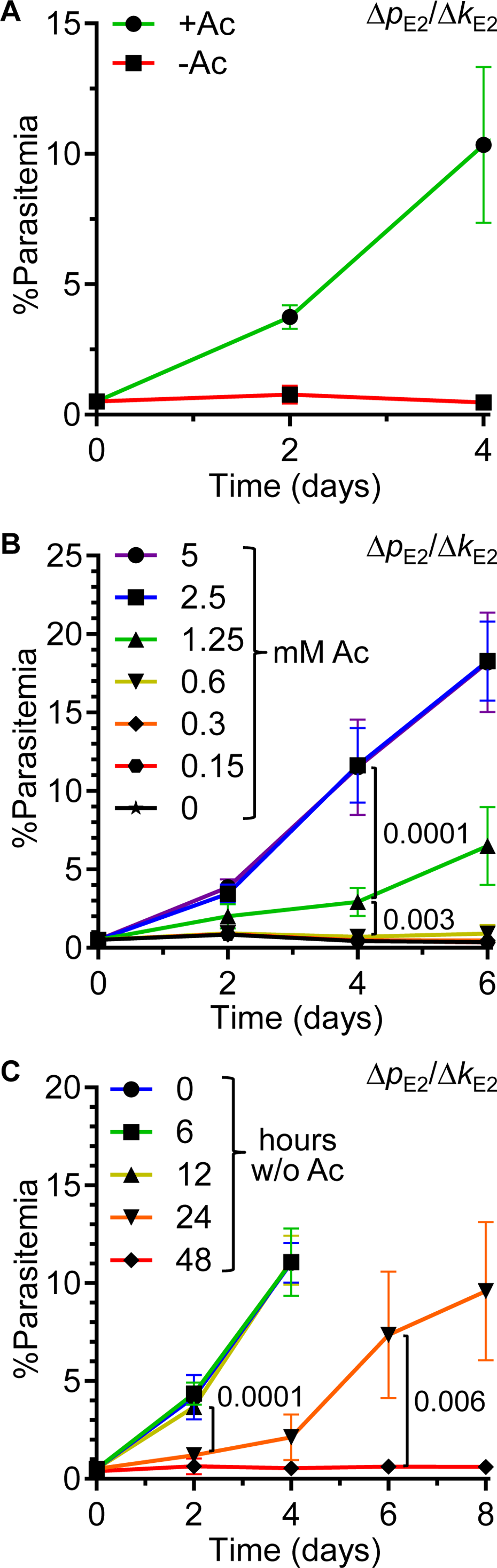
mPDH is essential in the absence of KDH. **A.** Growth curves comparing the growth of Δ*p*_E2_*/*Δ*k*_E2_ parasites in the presence and absence of 5 mM acetate (Ac). **B.** Growth of Δ*p*_E2_*/*Δ*k*_E2_ parasites in media supplemented with different concentrations of acetate (Ac). By day four of the growth curve, the parasitemia of the culture supplemented with 1.25 mM Ac (green) differed significantly from cultures supplemented with a two-fold higher (blue) or a two-fold lower (gold) concentration (two-way ANOVA, followed by Bonferroni’s correction; P values are provided in the plot). **C.** Survival of Δ*p*_E2_*/*Δ*k*_E2_ parasites after different periods of acetate (Ac) deprivation. Parasite growth assays were initiated after different periods of maintenance in acetate-free medium. The plots show the growth of these cultures after reintroduction of acetate. Significant parasite growth was observed on day two of the outgrowth period in the culture deprived for 12 hours relative to the culture deprived for 24 hours (two-way ANOVA, followed by Bonferroni’s correction; P values are provided in the plot). Error bars in the growth curves represent the standard deviation from at least two independent experiments, each conducted in quadruplicate.

Parasites survived but grew less well after 24 hours of acetate deprivation, and growth was not restored when acetate was added after 48 hours of deprivation (**Figure 3C**). Taken together, these results reveal a synthetic lethal phenotype between the *mpdh_E2_* and *kdh_E2_* genes presumably due to the capacity of both mPDH and KDH to generate acetyl-CoA.

### Mitochondrial acetyl-CoA biosynthesis is impaired in *Δp_E2_/Δk_E2_* parasites

Glucose and glutamine are the two major carbon sources for the TCA cycle and enter the mitochondrion as pyruvate and malate (from glucose) or α-ketoglutarate (from glutamine)^29^. To characterize the biochemical changes in the *Δp_E2_/Δk_E2_* parasite line, we used mass spectrometry-based metabolomics to compare the levels of acetyl-CoA and other TCA cycle intermediates in *Δp_E2_/Δk_E2_* and parental parasites by supplementing the growth media with isotopically labeled glucose (^13^C-glucose) and glutamine (^13^C-glutamine) (**Figure 4A**).

**Figure 4.**
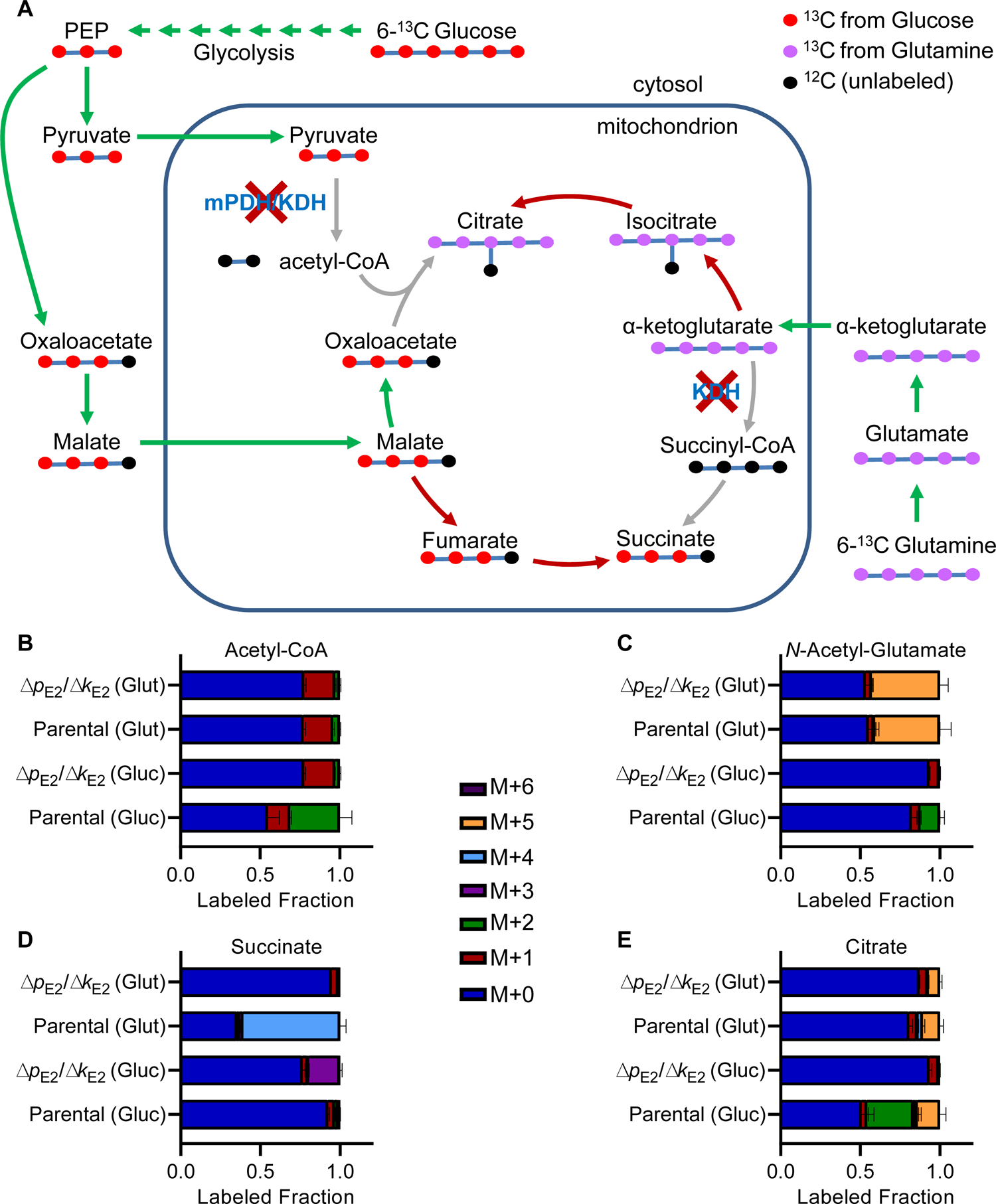
Mitochondrial acetyl-CoA biosynthesis is impaired in Δ*p*_E2_*/*Δ*k*_E2_ parasites. **A.** Schematic showing the labeling pattern of mitochondrial metabolites after short-term incubation with uniformly labeled ^13^C-glucose or ^13^C-glutamine. Red and purple circles represent the heavy isotope carbon atoms from glucose or glutamine, respectively. Black circles represent un-labeled atoms. Arrows represent enzymatic reactions and are colored based on whether labeled carbon atoms flow in the same direction as would be found in wild-type parasites (green), the reverse direction (red) or appear to be blocked due to the loss of mPDH and KDH activity. **B-E**. Fraction of isotopically labeled acetyl-CoA (**B**), *N*-acetyl glutamate (**C**), succinate (**D**), or citrate (**E**) when incubated with labeled glutamine (Glut) or glucose (Gluc) in parental or Δ*p_E2_/*Δ*k_E2_* parasites. For labeling experiments, color coding indicates the mass shift from the incorporation of heavy labeled carbon atoms in addition to the mass (M) of the parent compound. Labeling data are presented as the fraction of the total metabolite pool determined from N=3 experiments (parental) or N=2 experiments (Δ*p_E2_/*Δ*k_E2_*) with error bars representing the standard deviation (SD).

Supplementation with these carbon sources resulted in an average of over 85% of glucose-6-phosphate being derived from ^13^C-glucose, and over 60% of α-ketoglutarate being derived from glutamine across both lines for the duration of the labeling period (**Figure 4-figure supplement 1A,B**). We then assessed the incorporation of ^13^C label into acetyl-CoA from either glucose- or glutamine-derived carbon entering the mitochondrion. In parental parasites, acetyl-CoA and the acetyl-CoA metabolite N-acetyl-glutamate were labeled with two carbons from ^13^C-glucose, but this labeling was blocked in *Δp_E2_/Δk_E2_* parasites lacking mitochondrial PDH activity. ^13^C-glutamine did not label acetyl-CoA in both parental and *Δp_E2_/Δk_E2_* parasites, as expected (**Figure 4B**). By contrast, ^13^C-glutamine was readily incorporated into fully labeled *N*-acetyl-glutamate in both parasite lines, ruling out any inherent differences in biosynthesis of *N*-acetyl-glutamate between parental and *Δp_E2_/Δk_E2_* parasites (**Figure 4C**).

In addition to changes in acetyl-CoA production, we also observed perturbations to the TCA cycle in *Δp_E2_/Δk_E2_* parasites. In parental parasites, four carbons from glutamine were incorporated into succinate, whereas this process was blocked in *Δp_E2_/Δk_E2_* parasites, presumably because KDH was deleted in this line (**Figure 4D**). Similarly, carbons from ^13^C-glutamine were observed in fumarate and malate only in parental parasites (**Figure 4-figure supplement 1C,D**). The *Δp_E2_/Δk_E2_* parasites did, however, incorporate three carbons from labeled glucose into succinate, a process that was not observed in parental parasites (**Figure 4D**). This is likely a consequence of anaplerotic reactions that result in the import of malate into the mitochondrion (**Figure 4A**). Without KDH and mPDH activity, malate can be converted into oxaloacetate, but cannot progress further into citrate. Instead, excess malate results in the accumulation of fumarate and succinate using enzymes that normally function in the opposite direction^29^ (**Figure 4D**).

Analysis of citrate labeling provides further insight into TCA activity in *Δp_E2_/Δk_E2_* parasites (**Figure 4E**). In parental parasites treated with ^13^C-glucose, citrate was found to contain two labeled carbons (from acetyl-CoA) or five labeled carbons from the combination of acetate (two carbons) and oxaloacetate derived from anaplerotic malate (three carbons). Citrate did not contain this labeling pattern in *Δp_E2_/Δk_E2_* parasites, however, there appeared to be a small amount of five-carbon label from glutamine that may have been acquired through the reverse action of isocitrate dehydrogenase and aconitase (**Figure 4E**). Taken together, metabolomic analysis shows that the TCA cycle is blocked at two points in *Δp_E2_/Δk_E2_* parasites and generates products consistent with the failure of mPDH to make acetyl-CoA and the failure of KDH to make succinyl-CoA.

### KDH can support parasite growth in the absence of mPDH

Gene deletion and metabolomic studies in *P. falciparum* have provided strong evidence that KDH uses α-ketoglutarate as a substrate to produce succinyl-CoA^46^. Our results indicate that KDH can also produce acetyl-CoA in addition to its established role in the TCA cycle, but it is not clear whether KDH can produce sufficient acetyl-CoA in the mitochondrion to support normal parasite growth on its own. Individual deletion of mPDH_E1α_ and mPDH_E2_ indicates that mPDH is dispensable, however, it is possible that either mPDH_E1_ or mPDH_E2_ could function with KDH subunits in a heterologous complex, as observed in other organisms^50^. To assess the role of KDH alone, we generated a parasite line in which both mPDH subunits were deleted (*Δp_E1α_/Δp_E2_*) (**Figure 5-figure supplement 1A,B**). Growth assays of *Δp_E1α_/Δp_E2_* parasites showed that this line was able to grow without acetate supplementation (**Figure 5A**), suggesting that KDH can generate acetyl-CoA without the assistance of any mPDH subunits. To test this hypothesis, we used glucose and glutamine labeling, first confirming that *Δp_E1α_/Δp_E2_* parasites could convert glucose and glutamine into their initial metabolites (**Figure 5-figure supplement 1C, D**). We observed significantly decreased incorporation of ^13^C-glucose into acetyl-CoA (**Figure 5B**) and *N*-acetyl-glutamate (**Figure 5C**) at levels that were indistinguishable from those we observed for the *Δp_E2_/Δk_E2_* line (**Figure 4B,C**). Presumably, the production of acetyl-CoA by KDH in the *Δp_E1α_/Δp_E2_* line is sufficient for parasite growth, but is too low to be accurately measured by our methodology.

**Figure 5.**
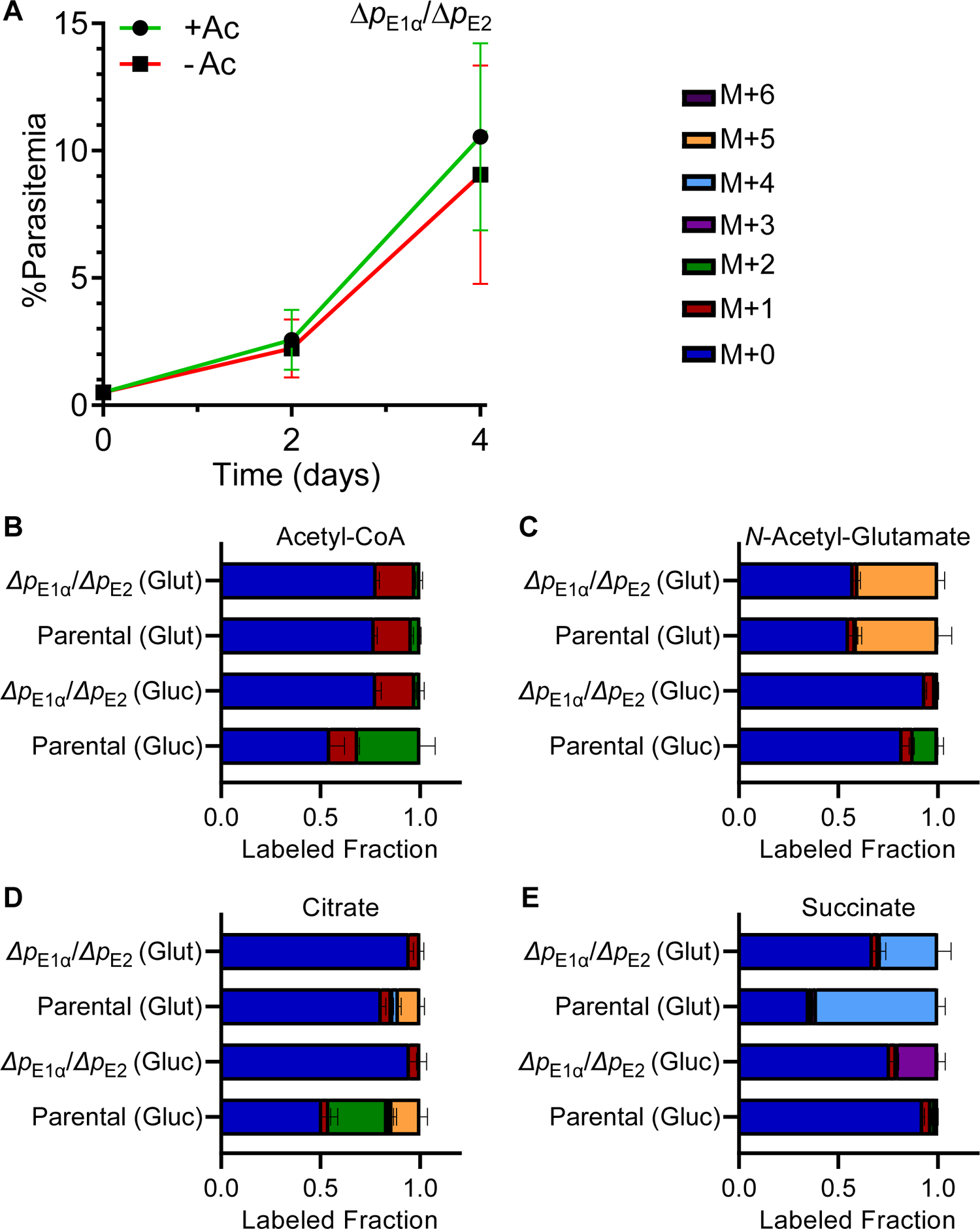
KDH can support parasite growth in the absence of mPDH. **A.** Growth curves comparing the growth of Δ*p*_E1α_*/*Δ*p*_E2_ parasites in the presence and absence of 5 mM acetate (Ac). Error bars represent the standard deviation from three independent experiments, each conducted in quadruplicate. **B-E.** Fraction of isotopically labeled acetyl-CoA **(B)**, *N*-acetyl-glutamate **(C)**, citrate **(D)**, or succinate **(E)** when incubated with labeled glutamine (Glut) or glucose (Gluc) in parental or Δ*p*_E1α_*/*Δ*p*_E2_ parasites. For labeling experiments, color coding indicates the mass shift from the incorporation of heavy labeled carbon atoms in addition to the mass (M) of the parent compound. Labeling data are presented as the fraction of the total metabolite pool determined from N=3 experiments (parental) or N=2 experiments (Δ*p*_E1α_*/*Δ*p*_E2_) with error bars representing the standard deviation (SD).

TCA cycle metabolites were perturbed in *Δp_E1α_/Δp_E2_* parasites in ways that closely paralleled the changes observed in the *Δp_E2_/Δk_E2_* parasite line. In both lines, carbon from glucose is not incorporated into citrate, consistent with the loss of acetyl-CoA synthesis from pyruvate (**Figure 5D**). Similarly, anaplerotic carbon (ultimately derived from ^13^C-glucose) accumulates in succinate in both lines (**Figure 5E**). In contrast to the *Δp_E2_/Δk_E2_* line, *Δp_E1α_/Δp_E2_* parasites could incorporate carbon from ^13^C-glutamine into succinate, demonstrating that KDH is still active in the mPDH deletion line (**Figure 5E**). Reduced levels of label incorporation (compared to the parental line) may be the result of altered TCA cycle flux. Indeed, carbon from glutamine does not seem to progress past succinate into fumarate and malate – the next metabolites in the TCA cycle (**Figure 5-figure supplement 1E,F**). In summary, these results show that *P. falciparum* malaria parasites can survive the loss of the mPDH complex, but there are significant impacts on acetyl-CoA production and other TCA cycle reactions.

### Protein lipoylation is required for acetyl-CoA biosynthesis

In malaria parasites, the attachment of lipoate to mitochondrial proteins proceeds through a unique two-step process. Lipoate is adenylated by an enzyme called LipL1 and then transferred to KDH_E2_ and mPDH_E2_ by an enzyme called LipL2 (**Figure 6A**)^41, 51^. Deletion of LipL2 should prevent KDH_E2_ and mPDH_E2_ lipoylation and mimic the phenotype of *Δp_E2_/Δk_E2_* parasites. As anticipated, LipL2 could only be deleted in the presence of acetate (**Figure 6-figure supplement 1A**) and the *Δlipl2* parasites were entirely dependent on high levels of acetate for growth (**Figure 6B, 6C**). We also found that the *Δlipl2* parasites could survive up to 24 hours of acetate deprivation, but growth was not restored when acetate was added after 48 hours of deprivation (**Figure 6D**). These results closely mirror those observed for the *Δp_E2_/Δk_E2_* line (**Figure 3**) and show that *Δlipl2* parasites phenocopy the *Δp_E2_/Δk_E2_* line. Deletion of LipL2 also results in the same metabolic perturbations previously observed in the *Δp_E2_/Δk_E2_* line. *Δlipl2* parasites could convert glucose and glutamine into their initial metabolites (**Figure 6-figure supplement 1B, C**), but could not incorporate ^13^C-glucose into acetyl-CoA (**Figure 6E**) or the downstream metabolite *N-*acetyl-glutamate (**Figure 6F**). The labeling of citrate (**Figure 6G**), succinate (**Figure 6H**), and other TCA cycle metabolites (**Figure 6-figure supplement 1D, E**) were also perturbed in the same manner as observed for the *Δp_E2_/Δk_E2_* line. Overall, these results are consistent with the conclusion that LipL2 is essential for the specific lipoylation of mPDH and KDH, and that loss of LipL2 results in loss of mPDH and KDH activity.

**Figure 6.**
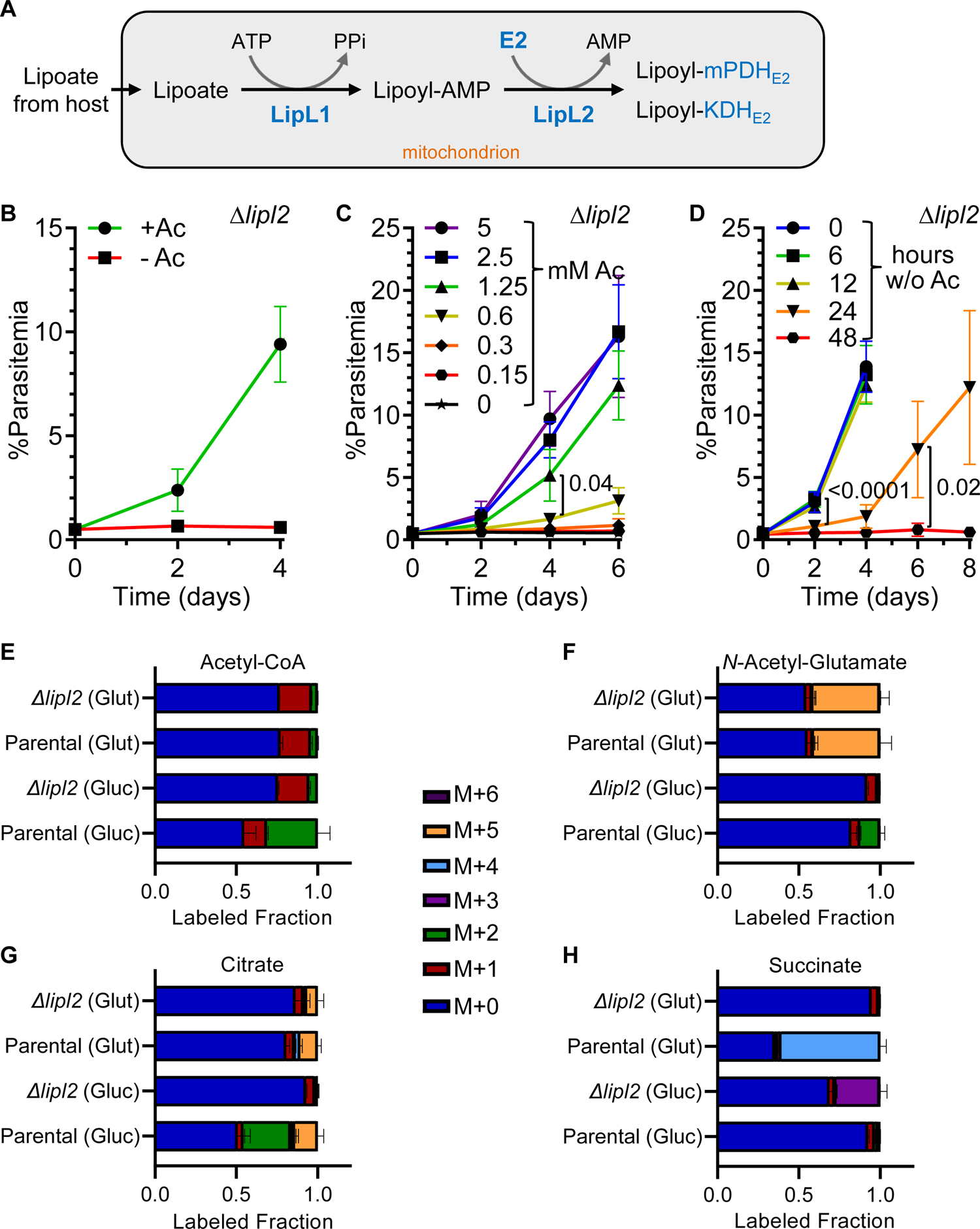
Lipoylation is essential for acetyl-CoA biosynthesis. **A.** Schematic showing the enzymatic steps required to attach lipoic acid (lipoate) to the E2 subunits of mPDH and KDH. Lipoate scavenged from the external environment is imported into the parasite mitochondrion and adenylated by lipoate ligase 1 (LipL1) in an ATP-dependent reaction. LipL2 transfers the lipoyl moiety from lipoyl-AMP to the E2 proteins. **B.** Growth curves comparing the growth of Δ*lipl2* parasites in the presence and absence of 5mM acetate (Ac). **C.** Growth of Δ*lipl2* parasites in media supplemented with different concentrations of acetate (Ac). By day four of the growth curve, the parasitemia of the culture supplemented with 1.25 mM Ac (green) differed significantly from the culture supplemented with a two-fold lower (gold) concentration (two-way ANOVA, followed by Bonferroni’s correction; P= 0.04). **D.** Survival of Δ*lipl2* parasites after different periods of acetate (Ac) deprivation. Parasite growth assays were initiated after different periods of maintenance in acetate-free medium. The plots show the growth of these cultures after reintroduction of acetate. Significant parasite growth was observed on day two of the outgrowth period in the culture deprived for 12 hours relative to the culture deprived for 24 hours (two-way ANOVA, followed by Bonferroni’s correction; P values are provided in the plot). **E-H.** Fraction of isotopically labeled acetyl-CoA **(E)**, *N*-acetyl-glutamate **(F)**, citrate **(G)**, or succinate **(H)** when incubated with labeled glutamine (Glut) or glucose (Gluc) in parental or Δ*lipl2* parasites. For labeling experiments, color coding indicates the mass shift from the incorporation of heavy labeled carbon atoms in addition to the mass (M) of the parent compound. Labeling data are presented as the fraction of the total metabolite pool determined from N=3 experiments (parental) or N=2 experiments (Δ*lipl2*) with error bars representing the standard deviation (SD). Error bars in the growth curves represent the standard deviation from at least two independent experiments, each conducted in quadruplicate.

### Acetate bypass of mitochondrial metabolism relies on nuclear Acetyl-CoA Synthetase

In *P. falciparum* parasites, labeled acetate is incorporated into acetyl-CoA, and the enzyme Acetyl-CoA Synthetase (ACS) has been hypothesized to catalyze this reaction^29, 32^. We explored this activity using *Δlipl2* and parental parasites labeled with ^13^C-acetate, ^13^C-glucose or ^13^C-glutamine. In contrast to previous labeling experiments, we maintained the parasites in medium containing 5 mM acetate during the glucose and glutamine labeling. We confirmed that labeled glucose and glutamine were converted into their initial metabolites (**Figure 7-figure supplement 1A, B**), but observed minimal incorporation of glucose carbon into acetyl-CoA (**Figure 7A**). This result suggested that the unlabeled acetate was incorporated into acetyl-CoA at the expense of labeled carbon from glucose. The labeling pattern in *N*-acetyl-glutamate supported this explanation since almost no glucose carbon was incorporated, while glutamine carbon was robustly incorporated in the glutamate moiety (**Figure 7B**). However, when we labeled parasite metabolites with 5 mM ^13^C-acetate, we observed incorporation of the label into the majority of the acetyl-CoA (M+2) and *N*-acetyl-glutamate over the course of the 2.5 hr labeling period (**Figure 7A, B**). Therefore, at very high concentrations acetate labeling is rapid and robust, even in the parental parasites with functional pathways to generate acetyl-CoA from glucose. Further metabolite analysis showed that supplemented acetate, even at high levels, is not incorporated into TCA cycle metabolites (**Figure 7-figure supplement 1B-E**), consistent with a lack of mitochondrial acetyl-CoA produced from ^13^C-acetate. Interestingly, ^13^C-acetate was incorporated into citrate, perhaps through a cytosolic reaction that remains to be characterized (**Figure 7-figure supplement 1F**).

**Figure 7.**
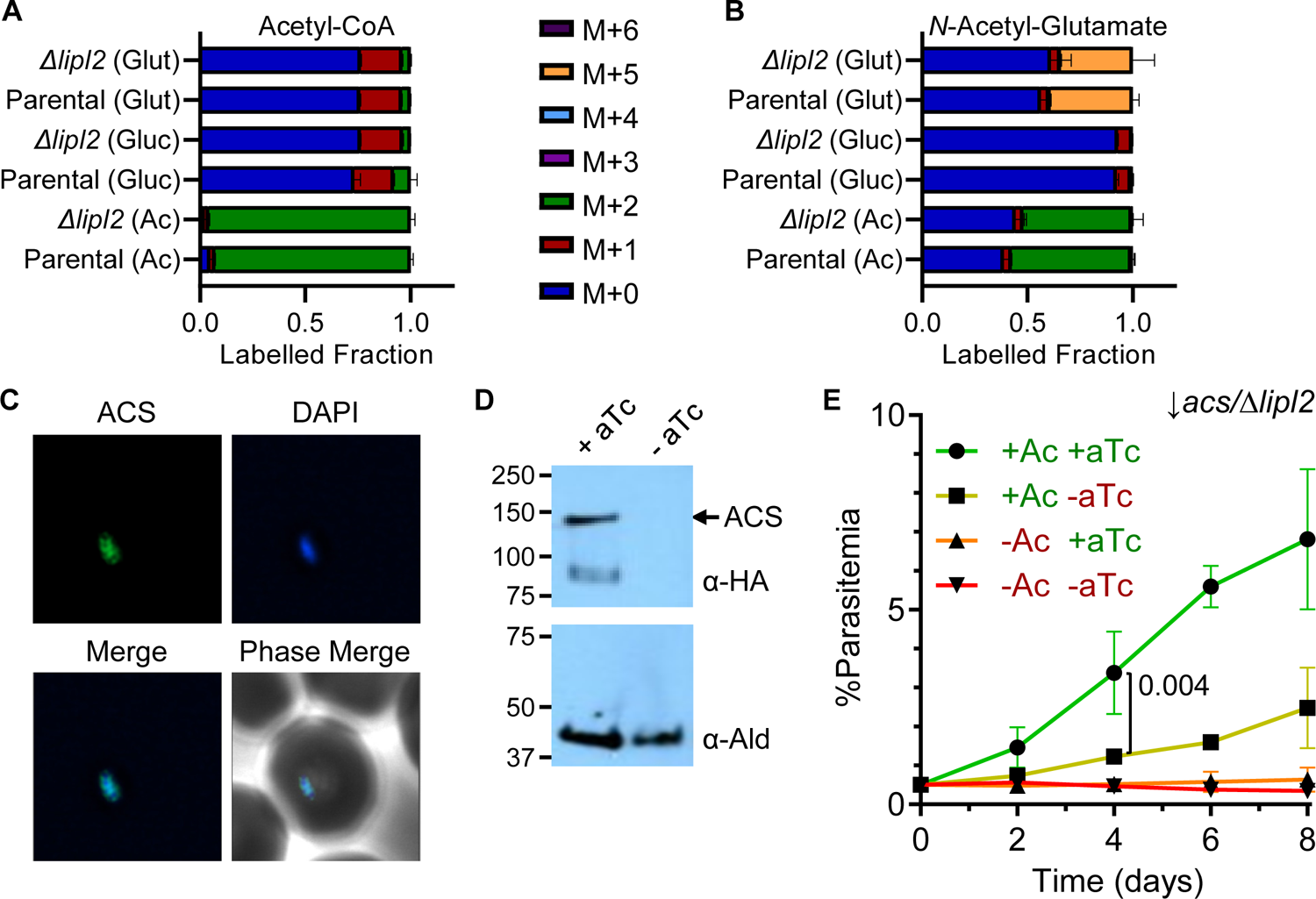
Acetyl-CoA biosynthesis using supplemented acetate is essential only in the absence of mitochondrial source. **A,B.** Fraction of isotopically labeled acetyl-CoA **(A)** or *N*-acetyl-glutamate **(B)** when incubated with labeled glutamine (Glut), glucose (Gluc) or acetate (Ac) in parental or Δ*lipl2* parasites. For labeling experiments, color coding indicates the mass shift from the incorporation of heavy labeled carbon atoms in addition to the mass (M) of the parent compound. Labeling data are presented as the fraction of the total metabolite pool determined from N=2 experiments with error bars representing the standard deviation (SD). **C.** Immunofluorescence co-localization of hemagglutinin-tagged ACS (green) with nuclear DNA stained with DAPI (blue) in fixed ↓*acs/Δlipl2* parasites. Images are 10 microns long by 10 microns wide. **D.** Western blot showing that ACS levels can be regulated with anhydrotetracycline (aTc) in the ↓*acs/Δlipl2* parasite line. After four days of aTc deprivation (-aTc), ACS-HA levels were below the limit of detection. Aldolase (Ald) served as a loading control. **E.** Growth curves of the ↓*acs/Δlipl2* parasite line grown under four different media conditions. Acetate was added (+Ac) in two of the conditions to bypass the loss of LipL2. Anhydrotetracycline was added (+aTc) in two of the conditions to maximize expression of ACS. In the presence of supplemented Ac, expression of ACS (+aTc) allowed parasites to outgrow ACS knockdown parasites (-aTc) by day 4 of the experiment (two-way ANOVA, followed by Bonferroni’s correction; P = 0.04). Error bars represent the standard deviation from two independent experiments, each conducted in quadruplicate.

Based on the metabolomics results (**Figure 7A)**, we hypothesized that the ACS enzyme must be responsible for almost all acetyl-CoA production in *ΔLipL2* parasites. To test this, we used a modified TetR-DOZI system^52^ plasmid pKD^53^ to knock down ACS expression in *Δlipl2* parasites, generating the *↓acs/Δlipl2* parasite line. Successful insertion of the pKD plasmid (**Figure 7-figure supplement 2**) resulted in tagging of ACS with 3xHA tag allowing us to localize the ACS protein to the parasite nucleus (**Figure 7C**) as has been recently reported^31–33^. We also confirmed that the knockdown was functional as we could no longer detect the ACS protein after parasites were cultured without aTc for four days (**Figure 7D**). We next tested the growth of the *↓acs/Δlipl2* line in acetate supplemented and non-supplemented conditions, while controlling the expression of the ACS enzyme using aTc. The *↓acs/Δlipl2* parasite line did not grow in the absence of acetate since there is no mitochondrial source of acetyl-CoA (**Figure 7E**). As expected, *↓acs/Δlipl2* parasites displayed a significant growth defect under non-permissive (no aTc) conditions when grown in 5 mM acetate (**Figure 7E**). These results show that ACS is responsible for the mobilization of scavenged acetate into acetyl-CoA and that this activity is essential in the absence of a mitochondrial source of acetyl-CoA.

### Acetyl-CoA produced in the mitochondrion is used throughout the parasite cell

Our data suggest that the mPDH and KDH enzymes in the mitochondrion are responsible for the production of acetyl-CoA, and that this activity is essential for the survival of blood stage parasites in the absence of acetate. These results raise the question of whether the acetyl-CoA produced in the mitochondrion is used throughout the cell for acetylation reactions. To address this question, we determined whether carbon from glucose can ultimately be incorporated into acetylated proteins found in other compartments of the cell (**Figure 8A**). We cultured NF54 strain parasites for four days in culture medium containing ^13^C-glucose and extracted parasite proteins for analysis by proteomic mass spectrometry. We observed the incorporation of ^13^C from glucose in several abundant acetylated proteins known to reside outside of the mitochondrion. In particular, we observed that nuclear histones were ^13^C-acetylated (**Table 2**), suggesting that mitochondrial acetyl-CoA is used in other compartments of the cell.

**Figure 8.**
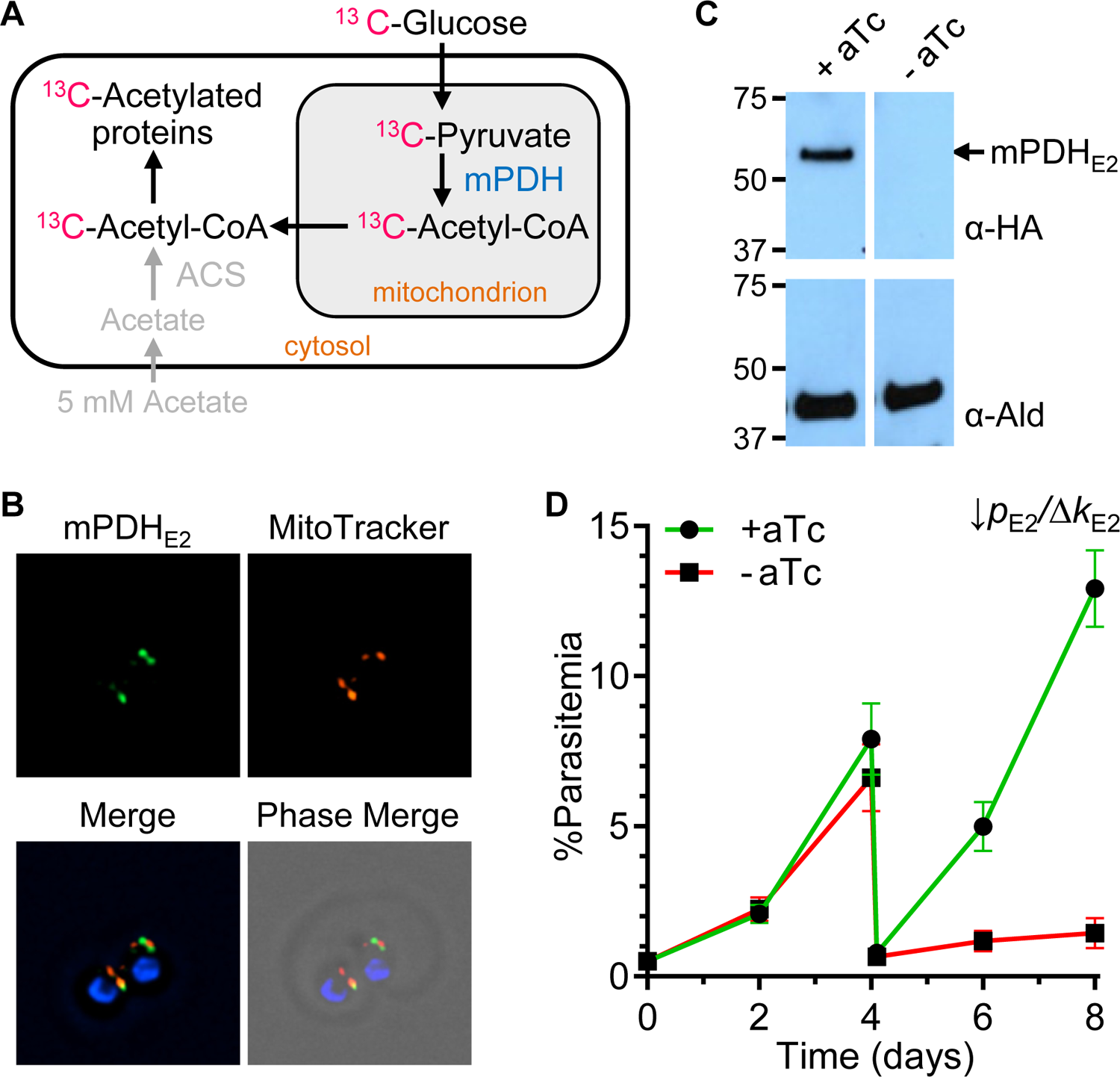
Mitochondrial acetyl-CoA is required for acetylation reactions in other cellular compartments. **A.** Illustration of the labeling experiment designed to determine whether acetyl-CoA synthesized in the mitochondrion is used for acetylation of cellular proteins. **B.** Immunofluorescence co-localization of hemagglutinin-tagged mPDH_E2_ (green) with MitoTracker (red) in fixed ↓*p*_E2_*/Δk*_E2_ parasites. Nuclear DNA was stained with DAPI (blue). Images are 10 microns long by 10 microns wide. **C.** Western blot showing that mPDH_E2_ levels can be regulated with aTc in the ↓*p*_E2_*/Δk*_E2_ line. After four days of aTc deprivation (-aTc), mPDH_E2_-HA levels were below the limit of detection. Aldolase (Ald) served as a loading control. **D.** Growth curves comparing the growth of ↓*p*_E2_*/Δk*_E2_ parasites in the presence and absence of anhydrotetracycline (aTc). After four days, the cultures were diluted by a factor of 10 to avoid overgrowth. Error bars represent the standard deviation from two independent experiments, each conducted in quadruplicate.

**Figure 9.**
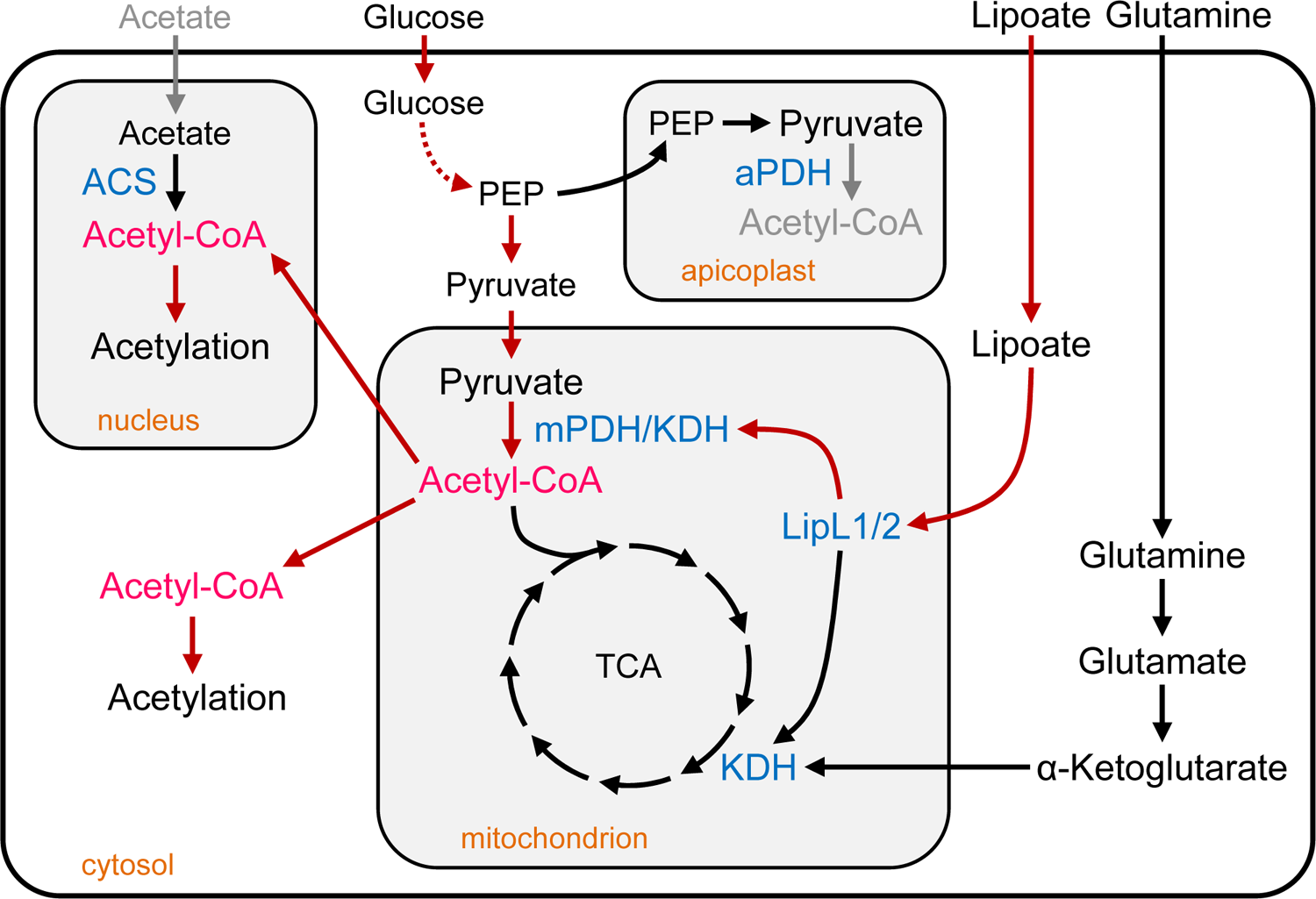
Metabolic model of acetyl-CoA synthesis in blood-stage *P. falciparum* **parasites.** Red arrows highlight processes required for the synthesis of acetyl-CoA and gray arrows indicate processes that are relevant to other stages of parasite development (aPDH activity in the apicoplast) or other cellular environments (scavenging of acetate). LipL1 and LipL2 require lipoate scavenged from the external environment to activate mPDH and KDH. Acetyl-CoA synthesized in the mitochondrion by mPDH and/or KDH is used to acetylate proteins in other compartments of the parasite. **Abbreviations:** LipL1, Lipoate Ligase 1; LipL2, Lipoate Ligase 2; PDH, pyruvate dehydrogenase; mPDH, mitochondrial PDH; aPDH, apicoplast PDH; KDH, ⍺-ketoglutarate dehydrogenase; ACS, acetyl-CoA synthetase; PEP, phosphoenolpyruvate; TCA, tricarboxylic acid cycle.

**Table 2.**
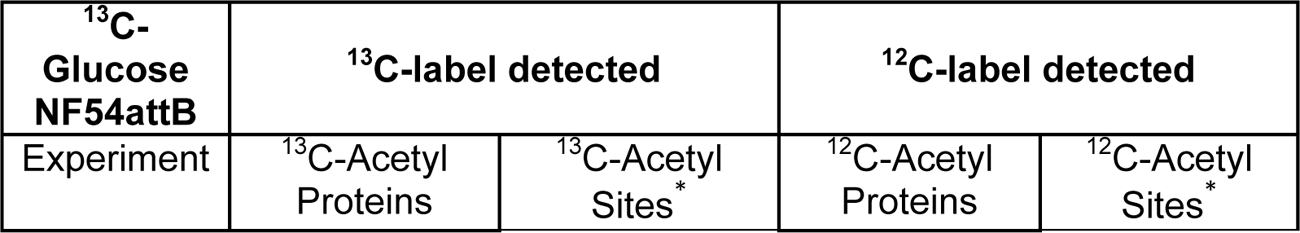
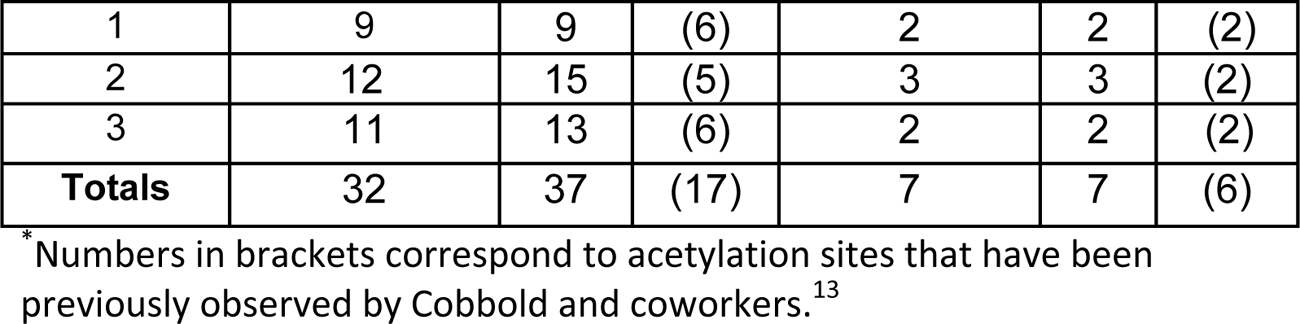
Incorporation of ^13^C-Glucose into protein acetylation sites in NF54 parasites.

| <sup>13</sup>C-Glucose NF54attB | <sup>13</sup>C-label detected |  | <sup>12</sup>C-label detected |  |
| --- | --- | --- | --- | --- |
| Experiment | <sup>13</sup>C-Acetyl Proteins | <sup>13</sup>C-Acetyl Sites* | <sup>12</sup>C-Acetyl Proteins | <sup>12</sup>C-Acetyl Sites* |

|  |  |  |  |  |  |  |
| --- | --- | --- | --- | --- | --- | --- |
| 1 | 9 | 9 | (6) | 2 | 2 | (2) |
| 2 | 12 | 15 | (5) | 3 | 3 | (2) |
| 3 | 11 | 13 | (6) | 2 | 2 | (2) |
| **Totals** | 32 | 37 | (17) | 7 | 7 | (6) |

\* Numbers in brackets correspond to acetylation sites that have been previously observed by Cobbold and coworkers.<sup>13</sup>

To confirm the link between the production of acetyl-CoA in the mitochondrion and protein acetylation throughout the cell, we generated a parasite line with conditional acetylation. Using the TetR-DOZI system plasmid pKD^53^ (described above) we generated an inducible mPDH_E2_ parasite line in which we also deleted the KDH_E2_. The resulting *↓p_E2_/Δk_E2_* line (**Figure 8-figure supplement 1**) should depend on the induction of mPDH (+aTc) to make acetyl-CoA in the mitochondrion and sustain growth. Detection of the 3xHA tag allowed us to localize the mPDH_E2_ protein to the mitochondrion (**Figure 8B**) and to confirm that the knockdown system was functional since we could no longer detect the mPDH_E2_ protein after parasites were cultured without aTc for four days (**Figure 8C**). Consequently, *↓p_E2_/Δk_E2_* parasites displayed a significant growth defect under non-permissive (no aTc) conditions compared to permissive (+ aTc) conditions after four days of growth (**Figure 8D**). Based on the knockdown and growth characteristics of this line, we conducted four-day ^13^C-labeling experiments. Parasites were split on the first day and cultured separately in permissive and non-permissive conditions. After two days of growth, the parasites were transferred into media containing ^13^C-glucose and cultured for the remaining two days of the experiment. Proteomic analysis of proteins extracted after the fourth day identified proteins acetylated with either ^12^C-acetate or ^13^C-acetate (**Table 3**). Under permissive conditions, the detected protein acetylation sites were predominantly labeled with carbon derived from ^13^C-glucose, consistent with the production of ^13^C-acetyl-CoA by mPDH in the mitochondrion. When mPDH was knocked down, ^12^C was observed in most acetylation sites, suggesting that mPDH activity was significantly reduced. Consistent with the direct ^13^C labeling experiments shown in **Table 2**, these results also show that acetyl-CoA produced in the mitochondrion by mPDH is essential for the acetylation of histones and other proteins located outside of the mitochondrion.

**Table 3.**
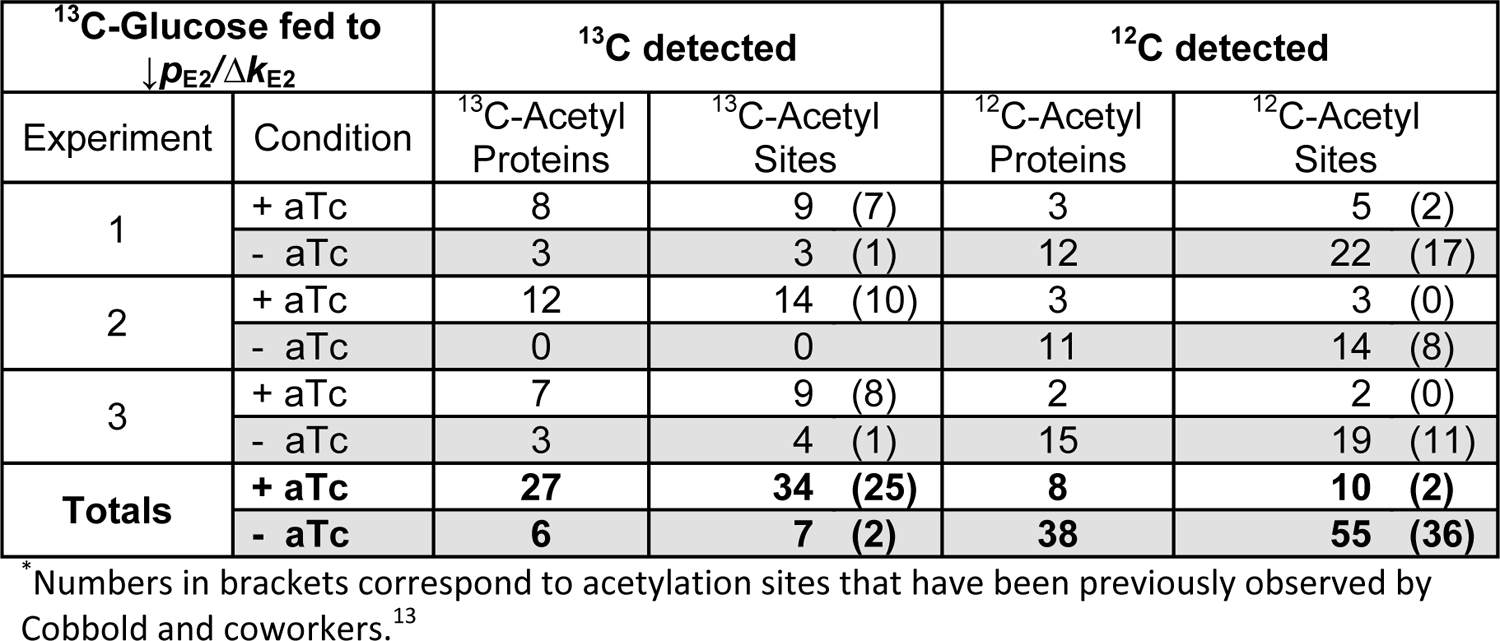
Incorporation of ^13^C-Glucose into protein acetylation sites in ↓*p*_E2_*/Δk*_E2_ parasites cultured under permissive (+aTc) or non-permissive (-aTc) conditions.

| $^{13}\text{C}$-Glucose fed to<br>$\downarrow p_{E2}/\Delta k_{E2}$ | | $^{13}\text{C}$ detected | | $^{12}\text{C}$ detected | |
| --- | --- | --- | --- | --- | --- |
| Experiment | Condition | $^{13}\text{C}$-Acetyl<br>Proteins | $^{13}\text{C}$-Acetyl<br>Sites | $^{12}\text{C}$-Acetyl<br>Proteins | $^{12}\text{C}$-Acetyl<br>Sites |
| 1 | + aTc | 8 | 9 (7) | 3 | 5 (2) |
|  | - aTc | 3 | 3 (1) | 12 | 22 (17) |
| 2 | + aTc | 12 | 14 (10) | 3 | 3 (0) |
|  | - aTc | 0 | 0 | 11 | 14 (8) |
| 3 | + aTc | 7 | 9 (8) | 2 | 2 (0) |
|  | - aTc | 3 | 4 (1) | 15 | 19 (11) |
| **Totals** | **+ aTc** | **27** | **34 (25)** | **8** | **10 (2)** |
|  | **- aTc** | **6** | **7 (2)** | **38** | **55 (36)** |

\*Numbers in brackets correspond to acetylation sites that have been previously observed by Cobbold and coworkers.<sup>13</sup>

## Discussion

The mitochondrion is typically thought of as the site of oxidative phosphorylation that provides energy for eukaryotic cells^54^. In blood-stage malaria parasites, components of the TCA cycle and the ATP synthase have been successfully deleted, demonstrating that oxidative phosphorylation is not required during this stage of parasite develpoment^46, 55^. What then are the essential roles of the mitochondrion in blood-stage parasites? The first answer to this question came from a seminal study showing that the mitochondrial electron transport chain is required for the oxidation of ubiquinone, which in turn is needed for the production of the pyrimidine orotate by the enzyme dihydroorotate dehydrogenase (DHOD)^56^. The antimalarial drug atovaquone potently inhibits the electron transport chain, but can be completely bypassed if the parasites are provided with an alternative mechanism to produce orotate, demonstrating that pyrimidines are essential products of the mitochondrion in blood-stage parasites^56^. These results led to the design of DHOD inhibitors to treat clinical malaria cases^57, 58^. A second essential product of the parasite mitochondrion is likely to be the precursor for iron-sulfur cluster synthesis in the cytosol by the cytosolic iron-sulfur cluster assembly (CIA) pathway^59^. Although there is no direct evidence in malaria parasites that the CIA pathway relies on the mitochondrion, this seems to be an aspect of mitochondrial metabolism that is extremely well conserved, even in organisms that contain highly reduced mitosomes and mitochondrion-like organelles^60, 61^. The results presented in the current work expose a third essential metabolite produced by the mitochondrion – acetyl-CoA. Our results show that acetyl-CoA is produced by ketoacid dehydrogenases in the mitochondrion of asexual blood stage *P. falciparum* parasites and is used for protein acetylation outside of the mitochondrion.

The enzyme complex originally annotated as BCKDH was an obvious candidate for mitochondrial PDH activity in *P. falciparum*. BCKDH (E2 subunit) was previously localized to the mitochondrion^41^ and was proposed to be the major source of acetyl-CoA when it was shown that the apicoplast PDH does not appreciably contribute to the acetyl-CoA pool or metabolic pathways downstream of this central precursor in blood-stage *P. falciparum* parasites^29^. A seminal study conducted in the murine malaria parasite *P. berghei* and the apicomplexan parasite *T. gondii* showed that deletion of the BCKDH E1α subunit significantly reduced the incorporation of carbon from glucose into TCA cycle metabolites and dramatically reduced parasite fitness^34^. Additionally, carbon from branched chain amino acids was not incorporated into acetyl-CoA, establishing the BCKDH as a mitochondrial PDH (mPDH) in these organisms^34^. Interestingly, the *P. berghei* E1α deletion line was no longer able to infect mature erythrocytes, but could only propagate in nutrient-rich reticulocytes^34^. This phenotype suggested that our *P. falciparum* mPDH E1α deletion line (*Δmpdh_E1α_*) would also fail to infect mature erythrocytes, but we did not observe a growth defect in this line or our *Δmpdh_E2_* line (**Figure 1E,F**). Instead, we found that the E2 subunits of mPDH and KDH form a synthetic lethal pair, indicating that both mPDH and KDH can produce acetyl-CoA in *P. falciparum* parasites (**Figure 3**). It may also be the case that KDH in *P. berghei* and/or *T. gondii* parasites could produce acetyl-CoA, albeit not at high enough levels to support normal growth.

In *P. falciparum* parasites, KDH participates in the TCA cycle and generates succinyl-CoA, which is then converted into succinate^46^. Deletion of the KDH E1 subunit blocked the incorporation of carbon from glutamine into succinate and led to the accumulation of KDH substrate α-ketoglutarate^46^. Thus, KDH functions as a TCA cycle enzyme in addition to producing acetyl-CoA. Dual specificity for KDH is consistent with biochemical experiments showing that both pyruvate and α-ketoglutarate are good substrates for recombinant *P. falciparum* KDH E1 with only a 2-fold difference in specific activity between the two substrates^49^. Presumably, the KDH E2 subunit would then function with the E1 subunit to produce acetyl-CoA from pyruvate. Alternatively, the mPDH and KDH could exchange subunits as has been reported previously in bacterial organisms^50^. To demonstrate that KDH alone can produce acetyl-CoA, we deleted both subunits of the mPDH and found that this line (*Δp_E1α/_Δp_E2_*) was able to grow without acetate supplementation (**Figure 5**). However, we did not observe significant incorporation of glucose carbon into acetyl-CoA in *Δp_E1α/_Δp_E2_* parasites and observed modest levels of label incorporation in other TCA cycle metabolites (**Figure 5, Figure 5-figure supplement 1**). Decreased catalytic efficiency of KDH (compared to mPDH) combined with active utilization of generated acetyl-CoA could explain why we did not detect labeled acetyl-CoA in *Δp_E1α/_Δp_E2_* parasites.

Our metabolite labeling experiments are consistent with previous studies showing that the TCA cycle is dispensable in blood-stage *P. falciparum* parasites^46^. The two main sources of TCA cycle carbon skeletons (glucose and glutamine) are completely blocked in the *Δpk_E2_* and *Δlipl2* lines (**Figure 4, Figure 6**). Unlike any of the TCA cycle deletion lines reported by Ke and coworkers^46^, our *Δpk_E2_* and *Δlipl2* lines require acetate supplementation for survival.

These results demonstrate that blood-stage parasites need acetyl-CoA produced in the mitochondrion for some purpose other than the TCA cycle. The data presented in **Table 2** and **Table 3** show that acetyl-CoA generated in the mitochondrion was used to acetylate both cytoplasmic and nuclear proteins, revealing an unexpected role for mitochondrial acetyl-CoA. This insight may extend to other malaria parasites, but does not apply to related parasites such as *T. gondii*. In *T. gondii*, multi-omic analysis of the BCKDH E1α deletion line shows that loss of mitochondrial acetyl-CoA results in few changes in the acetylome, transcriptome, and proteome in cellular compartments outside of the mitochondrion^62^. The acetyl-CoA pool in the nucleus and cytoplasm appears to be largely independent of the mitochondrion in *T. gondii* and relies on sources of acetyl-CoA that are not available to malaria parasites when they infect mature erythrocytes^62^. ATP citrate lyase (ACL) produces acetyl-CoA in the cytosol of *T. gondii*, but is not found in malaria parasites. The nuclear ACS enzyme in *T. gondii* may also produce more *de novo* acetyl-CoA in the nutrient-rich environment of nucleated host cells. ACS and ACL form a synthetic lethal pair^63^ in *T. gondii* and ablation of both enzymatic activities results in radical hypo-acetylation of histones, glycolytic enzymes and other nuclear/cytosolic proteins^62^.

Mitochondrial export of acetyl-CoA has been extensively studied in eukaryotic cells^64, 65^. One common method is to transport citrate formed from acetyl-CoA by citrate synthase to the cytosol where an ACL enzyme can then use citrate to regenerate acetyl-CoA^66^. Malaria parasites contain a mitochondrial citrate synthase^46^ and mitochondrial carrier proteins that could transport citrate^67, 68^, but do not seem to have an ACL enzyme, making the existence of a citrate shuttle unlikely. A second method of exporting acetyl-CoA is by forming acetyl-*L*-carnitine^69, 70^. However, a carnitine shuttle and the key mitochondrial enzyme carnitine acetyltransferase do not seem to exist in *Plasmodium* parasites. A third possibility involves the conversion of mitochondrial acetyl-CoA to acetate using a CoA transferase (with succinate as the other substrate) such as yeast Ach1^71^. Acetate released from the mitochondrion could be used to reform acetyl-CoA by the parasite ACS enzyme. A final possibility could be that an acetyl-CoA specific transporter is used to export acetyl-CoA from the mitochondrion even through this is not an activity typically found in mitochondria^72, 73^. An acetyl-CoA specific transporter identified in *Plasmodium* parasites was shown to be localized to the endoplasmic reticulum based on immunofluorescent staining^74^, and it seems unlikely that it would have a role in the mitochondrion as well. Alternatively, the carrier protein that is responsible for CoA import into the mitochondrion may help to export acetyl-CoA and this hypothesis could be tested once this carrier protein is identified. Although it remains to be determined whether acetyl-CoA is exported or a shuttle mechanism is used, our results show that acetyl-CoA generated in the mitochondrion is ultimately used elsewhere in the cell.

Our study also provides further insight into cofactor requirements during the asexual blood stages of *P. falciparum*. The KDH and mPDH enzyme complexes both require the cofactors thiamine pyrophosphate (TPP) and lipoate^35^. The thiamine antimetabolite oxythiamine kills blood-stage malaria parasites and dramatically reduces the ability of the parasites to incorporate carbon from glucose into acetyl-CoA^29, 49^. Although oxythiamine likely inhibits essential enzymes in isoprenoid biosynthesis^30^ and the pentose phosphate pathway^75^, its effect on acetyl-CoA synthesis can be explained through the inhibition of aPDH, mPDH, and KDH. The cofactor lipoate is also required for the activity of PDH and KDH enzymes^35^. Malaria parasites scavenge lipoate and covalently attach it to the E2 subunits of mPDH and KDH as well as a third mitochondrial protein called the H-protein^40^. Biochemical studies explained how lipoate could be attached to these three mitochondrial proteins in malaria parasites.

Recombinant lipoate ligase (LipL1) catalyzes the formation of lipoy-AMP from lipoate and ATP and can subsequently transfer the lipoyl moiety to the H-protein, but it does not recognize KDH or mPDH as substrates^41^. A lipoyl transferase called LipL2 is required to attach lipoate to the E2 subunits of KDH and mPDH using lipoyl-AMP made by LipL1^51^. If LipL2 has the same enzymatic activity in the complex environment of the parasite mitochondrion, we would expect deletion of LipL2 to result in the inactivation of both KDH and mPDH. The growth, acetate dependence, and metabolism of *Δlipl2* parasites closely mimics that of the double deletion line (*Δp_E2_/Δk_E2_*), confirming this hypothesis (**Figure 6**). The dependence of the *Δlipl2* line on acetate supplementation further suggests that LipL2 does not have any other essential roles in parasite biology other than supporting the generation of acetyl-CoA.

## Conclusions

In this report, we show that the mitochondrion of malaria parasites is the primary source of the essential metabolite acetyl-CoA, and that this metabolite is exported from the mitochondrion for use in other compartments of the cell. Surprisingly, two enzymes are capable of synthesizing acetyl-CoA in the mitochondrion and both must be inactivated to halt the growth of blood-stage parasites. One route to inactivate both enzymes is by preventing the attachment of the cofactor lipoate catalyzed by LipL2. Deletion of LipL2 blocks acetyl-CoA synthesis in the mitochondrion and results in parasites that can only survive when supplemented with high levels of acetate in their growth medium. We show that the ACS enzyme is responsible for the survival of parasite lines deficient in mitochondrial acetyl-CoA synthesis by mobilizing exogenous acetate to form acetyl-CoA. However, this metabolic bypass is unlikely to support parasite growth *in vivo* since more than 1 mM acetate is required, which is substantially higher than the concentration that has been reported in human blood^76^. Ultimately, this work links the scavenging of the host metabolite lipoate to the production of an essential mitochondrial metabolite which then can alter a broad range of cellular processes through the acetylation of proteins including histones, transcription factors and metabolic enzymes. The prospect of a mitochondrial pathway controlling many of these critical cellular processes supports the further exploration of mitochondrial acetyl-CoA biosynthesis as a novel pathway for antimalarial therapeutic interventions.

## Materials and Methods

### *P. falciparum* culture and maintenance

Unless otherwise noted, blood-stage *P. falciparum* parasites were cultured in human erythrocytes at 1% hematocrit in CMA (Complete Medium with Albumax) containing RPMI 1640 medium with L-glutamine (US Biological Life Sciences), supplemented with 20 mM HEPES, 0.2% sodium bicarbonate, 12.5 μg/mL hypoxanthine, 5 g/L Albumax II (Life Technologies) and 25 μg/mL gentamicin. Cultures were maintained in 25 cm^2^ gassed flasks (94% N_2_, 3% O_2_, 3% CO_2_) and incubated at 37 °C. For cultures that require acetate supplementation, 5 mM sterile sodium acetate was supplemented in the media.

### Generation of knockout and knockdown constructs

We employed Cas9-mediated gene editing^77^ using plasmid pCasG-LacZ^53^ to express Cas9 and a gRNA and repair plasmid pRSng^30, 78^. For generation of pRSng, homology arms of ∼200-400 bp from the gene of interest were amplified using homology arm (HA) 1 and 2 forward and reverse primers (**supplementary file 1)** using blood stage *P. falciparum* NF54^attB^ genomic DNA as template. HA1 and HA2 primers were designed to contain ∼15 bp overhangs for insertion into pRSng using ligation-independent cloning (LIC) methods (In-Fusion, Clontech). pRSng plasmid was digested with *Not*I for insertion of HA1, and with *Ngo*MIV for HA2. Plasmid pCasG-LacZ was digested with *Bsa*I for insertion of double-stranded DNA adaptamers (generated from primers listed in **supplementary file 1**) using LIC or standard ligation; positive colonies were selected using blue/white colony screening. To generate double knockout lines, the gene encoding human Dihydrofolate Reductase (*dhfr*) was excised from plasmid pRSng using *Bam*HI/*Hin*dIII and replaced using LIC (primers pRsBSD.F and pRsBSD.R) with a sequence encoding *Aspergillus terreus* Blasticidin-S Deaminase codon harmonized for expression in *P. falciparum*^79^ to generate the pRSngB plasmid.

For knockdown constructs, we used a variant of plasmid pMG74^52^ called pKD^53^. For generation of the *↓mpdh_E2_* (mPDH_E2_ knockdown) plasmid, HA1 (253 bp) was amplified from bases 1592 to 1844 of the sequence encoding mPDH_E2_. HA2 (327 bp) was amplified from the 3’ UTR, starting from 471 bp after the stop codon. The HA2 and HA1 amplicons were concatenated by PCR using the HA2 forward primer and the HA1 reverse primer to generate a HA2-HA1 amplicon with *Eco*RV endonuclease sites between the two homology arms. The HA2-HA1 fragment was then inserted into pKD digested with *Asc*I and *Aat*II. The sequences of primers used for amplifying homology arms and the gRNA sequences are included in **supplementary file 1**. For generation of the *↓acs* (ACS knockdown) plasmid, HA1 (323 bp) was amplified from bases 3169 to 3491 of the sequence encoding ACS. HA2 (247 bp) was amplified from the 3’ UTR region starting 65 bp after the stop codon. HA2 and HA1 were concatenated and inserted into the knockdown plasmid as described above for mPDH_E2_. The guide RNA was cloned into the pCasG-LacZ vector as described above.

### ***P.*** *falciparum* transfections to generate knockout and knockdown lines

For deletion of *mpdh_E1_*_α_ (PF3D7_1312600), *mpdh_E2_* (PF3D7_0303700), *e3* (PF3D7_1232200), *kdh_E2_* (PF3D7_1320800) and *lipl2* (PF3D7_0923600), 300 μL of red blood cells were electroporated with 75 μg each of gene-specific pRSng and pCasG plasmids.

Electroporated RBCs were mixed with ∼2.5 mL NF54^attB^ parasites synchronized as schizonts and cultured in CMA (or CMA with 5 mM acetate for some experiments). After 48 hrs, transfected parasites were selected with 1.5 μM DSM1 and 2.5 nM WR99210 for seven days, after which they were cultured in drug-free medium until parasites were observed (typically between 17 and 30 days after transfection). Once parasites were observed, they were switched to medium containing WR99210. For generation of double knockout parasites, we used the pRSngB plasmid containing *mpdh_E2_* homology arms. The *Δkdh_E2_* and *Δmpdh_E1α_* parasite lines were transfected with the pRSngB plasmid using the methods described above with WR99210 substituted with 2.5 μg/mL blasticidin. After parasites were observed, they were maintained in media containing WR99210 and blasticidin.

For the generation of *acs* (PF3D7_0627800) and *mpdh_E2_* (PF3D7_1314600) inducible knockdown lines, plasmid pKD was linearized by overnight digestion with *Eco*RV. Transfection was performed by electroporating 75 μg of linearized pKD along with 75 μg of the corresponding pCasG plasmid into 300 μL of uninfected red blood cells. Following electroporation, 2.5 mL of NF54^attB^ parasites were added to red blood cell cultures containing the plasmids. Starting 48 hrs post transfection, parasites were maintained in drug selection media containing 1.5 μM DSM1 and 2.5 μg/mL blasticidin for 7 days, after which drug-free media was used until parasites were observed (20-30 days post-transfection). After this point, parasites were maintained in medium containing blasticidin.

### Confirmation of genotypes

Gene deletion parasite lines generated with the pRSng plasmid were validated with genotype PCR reactions. Primers were designed to screen for 5’ integration (Δ5’ reaction primers GOI.5’F and pRS.R) and 3’ integration (Δ3’ reaction primers pRS.F and GOI.3’R) of the gene of interest (GOI) disruption cassette, and the 5’ region (primers GOI.5’F and GOI.5’WTR) and 3’ region (primers GOI.3’WTF and GOI.3’R) of the unmodified gene. The parental NF54^attB^ line was used as a control for these reactions. For the ACS knockdown line, primers were designed to screen for 5’ integration (Δ5’ reaction primers iACS.5’F and pMG.3HA.R) and 3’ integration (Δ3’ reaction primers HSP86.F and iACS.3’R) of the pKD plasmid, and the unmodified parental gene (P reaction primers iACS.5’F and iACS.3’R). For the mPDH_E2_ knockdown line, primers were designed to screen for 5’ integration (Δ5’ reaction primers imPDH.5’F and pMG.3HA.R) and 3’ integration (Δ3’ reaction primers HSP86.F and imPDH.3’R) of the pKD plasmid, and the unmodified parental gene (P reaction primers imPDH.5’F and imPDH.3’R). The parental NF54^attB^ line was used as a control for these reactions. The primer sequences are included in **supplementary file 1**.

### Parasite growth curve determination

For growth curve determination, parasites were stained with SYBR Green, and analyzed via flow cytometry. Parasite lines of interest were each seeded in quadruplicates in 96-well plates at about 0.5% parasitemia. Parasitemia was then measured at two-day intervals as previously described^78^. Parasites were diluted 1:10 in CMA and stored in a 96-well plate at 4 °C until analysis by flow cytometry. Parasites were stained by transferring 10 µL of the 1:10 dilutions to a 96-well plate containing 100 µL of 1x SYBR Green (S7563, Invitrogen) in PBS, and incubated for 30 minutes while shaking. Post-incubation, 150 µL of PBS was added to each well to dilute unbound SYBR Green dye. Samples were analyzed with an Attune Nxt Flow Cytometer (Thermo Fisher Scientific), with a 50 µL acquisition volume, and a running speed of 25 µL/minute with 10,000 total events collected.

### Immuno-fluorescent microscopy

Parasites were resuspended in a 1:2 solution of 4% paraformaldehyde and 0.0075% glutaraldehyde in PBS for 30 minutes at 37° C to fix cells. Next, 40 µL of fixed parasite sample was added to 11 mm wells in PTFE printed microscope slides (Electron Microscopy Sciences, Cat No: 63422-11) and incubated for half an hour. After 30 minutes, superficial fluid was aspirated and wells were washed once with 40 µL of PBS. Cells were then permeabilized for 10 minutes with 40 µL of 1% Triton X-100 solution followed by three washes with PBS. Cells were then reduced for 10 minutes by adding 40 µL of 0.1 g/L NaBH_4_ in PBS and washed 3 times with PBS afterwards. Cells were blocked for 2 hrs in a solution of 3% (30 g/L) BSA in PBS, and then washed 3 times with PBS. Cells were then incubated with 40 µL of primary antibody (1:1,000 rat anti-HA mAb 3F10, Roche) at 4°C overnight in a solution containing 3% BSA in PBS. Cells were washed three times with PBS on the next day, and then incubated for 2 hrs in 40 µL of secondary antibody (1:1,000 anti-rat Alexa Fluor 488, Life Technologies) in 3% BSA in PBS. Cells were washed with PBS 3 times and stained with Gold DAPI antifade (Life Technologies) under a coverslip sealed with nail polish. Slides were then viewed using a Zeiss AxioImager M2 microscope. A series of images spanning 5 µm were acquired with 0.2 µm spacing and images were deconvolved with VOLOCITY software (PerkinElmer) to report a single combined z-stack image.

### Western blot

For western blot analysis, 10 mL of parasite culture of about 10-12% parasitemia in 2% hematocrit was pelleted by centrifugation. Pelleted cells were saponin lysed in 0.1% saponin for 5 minutes on ice. Lysed parasites were pelleted and washed once to remove soluble red blood cell contents. The parasite pellet was then resuspended in 50 µL NuPAGE LDS sample buffer (Thermo Fisher) and boiled and vortexed for 10 minutes. Proteins were resolved by SDS-PAGE on 4-12% gradient gels and transferred to nitrocellulose membranes. The membranes were then blocked for an hour in 5% milk powder/PBS solution and probed with 1:2,500 rat anti-HA mAb 3F10 antibody (Roche) in 5% milk powder/PBS solution overnight. After washing in PBS containing 0.5% Tween-20, the membranes were probed with 1:2,500 goat anti-rat HRP-linked secondary antibodies (GE healthcare, NA935) in 5% milk powder/PBS solution for 1 hour, and then washed again and the signal detected using Super Signal Chemiluminescent substrate (Thermo fisher scientific, 34577). For loading controls, the membranes were stripped in 200 mM Glycine, 1% Tween-20 for 15 min and washed in PBS containing 0.5% Tween-20. Aldolase was detected using the above protocol with 1:10,000 mouse anti-aldolase antiserum (a kind gift from David Sullivan) and 1:2,500 sheep anti-mouse HRP-linked secondary antibodies (GE healthcare, NA931).

### 13C-Metabolic labeling experiments to detect acetylated proteins

Blood stage parasite cultures were initiated in 50 mL culture flasks at 2% hematocrit, at ∼1% parasitemia and grown for four days. Wild type parasites were grown in glucose-free media that was supplemented with 2 g/L of ^13^C-glucose for all four days. For the knockdown experiments, the *↓mpdh_E2_/Δkdh_E2_* line (*↓p_E2_/Δk_E2_*) was grown in regular media for first two days and media with ^13^C-glucose for the last two days in permissive (0.5 µM aTc) or non-permissive (No aTc) conditions. After 4 days of growth, cells were harvested through centrifugation at 2,000 g for 10 minutes. The cell pellet was resuspended in 1 mL of 0.05% saponin and incubated on ice for 5 minutes to selectively lyse red blood cells. After incubation, the lysis solution was diluted with 10 mL of PBS and centrifuged at 2,000 g for 10 minutes. Pelleted parasites were washed once with PBS to remove red blood cell residue. Parasite pellets were then resuspended in 1 mL of PBS and spun down again at 16,000 g for 5 minutes. Protein extraction was performed by resuspending the pellet in 50µL of guanidine-based extraction buffer (8 M guanidine HCl, 100 mM Tris, pH 8.0, 0.5 mM DTT), briefly vortexing, and incubating the solution on ice for 20-30 minutes. After the incubation, the solution was spun down and the supernatant containing extracted protein was stored at −80 °C for mass spectrometric analysis. The protein sample was quantified using Bio-Rad Protein Assay Dye Reagent (Cat No: 5000006).

### Mass spectrometry (LC-MS/MS) based identification of acetylated peptides

Protein extracts (∼20 µg) were reduced with 20 µL of 150 mM dithiothreitol for 45 mins at 60 °C, and subsequently alkylated with 20 µL of 250 mM iodoacetamide in dark for 15 minutes. Samples were digested by adding 100 µL trypsin solution (10 mM TEAB, pH 7.5, 2 µg Promega trypsin #V5111) and incubated overnight at 37 °C. Peptides were acidified and buffer exchanged using the Oasis Plate HLB (Waters) collected and dried by speedvac. Peptide fractions were analyzed by liquid chromatography interfaced with tandem mass spectrometry (LC/MSMS) using an Easy-LC 1100 HPLC system (Thermofisher) interfaced with an Orbitrap Fusion Lumos Mass Spectrometer (Thermofisher). Fractions were resuspended in 20 µL loading buffer (2% acetonitrile in 0.1% formic acid) and analyzed by reverse phase liquid chromatography coupled to tandem mass spectrometry. Peptides (20% each fraction) were loaded onto a C18 trap (S-10 µM, 120 Å, 75 µm x 2 cm, YMC, Japan) and subsequently separated on an in-house packed PicoFrit column (75 µm x 200 mm, 15 µm, +/-1 µm tip, New Objective) with C18 phase (ReproSil-Pur C18-AQ, 3 µm, 120 Å, Dr.Maisch, Germany) using 2-90% acetonitrile gradient at 300 nL/min over 120 min. Eluting peptides were sprayed at 2.0 kV directly into the Lumos. Survey scans (full MS) were acquired from 370-1400 m/z with data dependent monitoring with a 3 sec cycle time. Each precursor was individually isolated in a 1.2 Da window and fragmented using HCD activation collision energy 30 and 15 sec dynamic exclusion, first mass being 120 m/z. Precursor and fragment ions were analyzed at resolutions of 120,000 and 30,000, respectively, with automatic gain control (AGC) target values at 4e5 with 60 ms maximum injection time (IT) and 1e5 with 100 ms maximum IT respectively. Isotopically resolved masses in precursor (MS) and fragmentation (MS/MS) spectra were processed in Proteome Discoverer (PD) software (v2.4, Thermo Scientific). All MS/MS samples were analyzed using Mascot (Matrix Science, London, UK; version 2.6.2). Mascot was set up to search the UP000001450_36329_3D7.fasta; RefSeq_83_Human_170919 database (118,734 entries) assuming the digestion enzyme trypsin. Mascot was searched with a fragment ion mass tolerance of 0.015 Da and a parent ion tolerance of 3.0 PPM. Carbamidomethyl of cysteine was specified in Mascot as a fixed modification. Deamidated products of asparagine and glutamine, oxidated methionine, and acetylated lysine (with 0, 1, or 2 ^13^C-labeled carbons) were specified in Mascot as variable modifications. Scaffold (version Scaffold_4.11.0, Proteome Software Inc., Portland, OR) was used to validate MS/MS-based peptide and protein identifications. Peptide identifications were accepted if they could be established at greater than 95.0% probability by the Peptide Prophet algorithm^80^ with Scaffold delta-mass correction. Protein identifications were accepted if they could be established at greater than 95.0% probability and contained at least 1 identified peptide. Protein probabilities were assigned by the Protein Prophet algorithm^81^. Proteins that contained similar peptides and could not be differentiated based on MS/MS analysis alone were grouped to satisfy the principles of parsimony.

### Whole genome sequencing of the clonal lines

Parasites used for gDNA extraction were grown to 8-10% schizonts in a 50-mL culture at 4% hematocrit on the day of extraction. Cells were isolated by centrifugation (1,500 rpm for 3 minutes) before aspirating the supernatant and adding 50 mL of 0.1% saponin in PBS. Following 10 minutes of lysis, parasites were isolated by centrifugation (3,000 rpm for 10 minutes) and washed twice with 50 mL of PBS. The gDNA was extracted using a Qiagen DNeasy® kit according to provided protocol. Extracted gDNA was quantified using a Nanodrop spectrophotometer. For sequencing, 1 µg of gDNA per sample was analyzed using the TruSeq® DNA PCR-Free Low-Throughput Library Prep Kit (Illumina) according to the manufacturer’s directions. Genomic sequence data for all genetically-modified lines are available at the NCBI Sequence Read Archive under submission SUB9847689 (https://account.ncbi.nlm.nih.gov/?back_url=https%3A//submit.ncbi.nlm.nih.gov/subs/sra/SUB9847689/).

### 13C-Metabolic labeling experiments to detect small metabolites

*Plasmodium falciparum* parasites used for metabolomics experiments were tightly synchronized one cycle prior to performing the metabolite extraction procedure. On the day of metabolite extraction, parasites were cultured to 5-10% early trophozoites at 4% hematocrit. Parasite lines requiring acetate for survival were grown in presence of 5 mM acetate. Parasites were then magnetically purified from uninfected RBCs and resulting culture was consolidated in fresh media without additional acetate. Parasite concentration was determined using a hemocytometer and adjusted to 1×10^8^ cells/mL. For each labeling experiment, 1 mL of parasites was added to 4 mL of media in triplicate per each condition. The media used for labeling was deprived of labeled metabolite: glucose, glutamine, or acetate. Before initiating incubation, ^13^C_6_-glucose, ^13^C_5_-glutamine, or ^13^C_2_-acetate were added to a concentration equivalent to standard culture conditions (11 mM for glucose, 2 mM for glutamine, and 5 mM for acetate). Parasites were incubated under these conditions for 2.5 hrs. The labeling media were then aspirated, and the parasite pellets were washed with PBS followed by quenching with 1 mL of 90% methanol containing 0.25 µM ^13^C_4_,^15^N_1_-aspartate. These samples were vortexed for 30 seconds and centrifuged for 15 minutes at 14,000 rpm at 4 °C. Following centrifugation, these samples were dried under nitrogen gas and stored at −80 °C. To prepare these samples for HPLC/MS analysis, they were removed from storage and placed on ice for 10 minutes to thaw. Samples were resuspended in 100 µL of 3% HPLC-grade methanol:water containing 1 µM chlorpropamide to achieve a final concentration of 1×10^9^ cells/mL in each sample. Samples were centrifuged and transferred to vials for HPLC/MS analysis. An equivalent portion of all samples was pooled into a separate vial to serve as a quality control for metabolite detection.

Samples were run using a Thermo Scientific Exactive Plus mass spectrometer. Chromatographic conditions were similar to those used previously^6^. Briefly, the liquid chromatography column used was a Waters Xselect C18 HSS T3 column with 2.5 µm particle diameter. The mobile phase was composed of 3% methanol with 10 mM tributylamine, 2.5 µM medronic acid, and 15 mM acetic acid (solvent A) and methanol (solvent B). The mass spectrometer was operated exclusively in negative mode with a scan range from 85-1000 m/z. Data analysis was performed using El-Maven mass spectrometry analysis software^82^.

Processed data for the metabolites presented in this manuscript are available in **supplementary file 2**. Raw data for all metabolites are available at the NIH workbench under submission ST002024 (http://dev.metabolomicsworkbench.org:22222/data/DRCCMetadata.php?Mode=Study&StudyID=ST002024).

## Acknowledgements

We thank David Sullivan for his kind gift of mouse anti-aldolase antiserum. This work was supported by the National Institutes of Health R01 AI125534 (STP), the Johns Hopkins Malaria Research Institute, and the Bloomberg Family Foundation. ML was supported by the Eberly College of Science and the Huck Institutes of the Life Sciences at The Pennsylvania State University. JTM was supported by training grant T32 DK120509. The funders had no role in study design, data collection and analysis, decision to publish, or preparation of the manuscript. We also thank the Huck Institutes of Life Sciences Metabolomics Core Facility at Penn State University.

## Author contributions

SCN, JTM, ML and STP arranged the figures and wrote the manuscript, with input from all listed coauthors. SCN, JTM and AM carried out the experiments displayed in this manuscript.

## Conflict of interests

The authors declare that they have no conflict of interest.

---

**Figure 1-source data 1.**
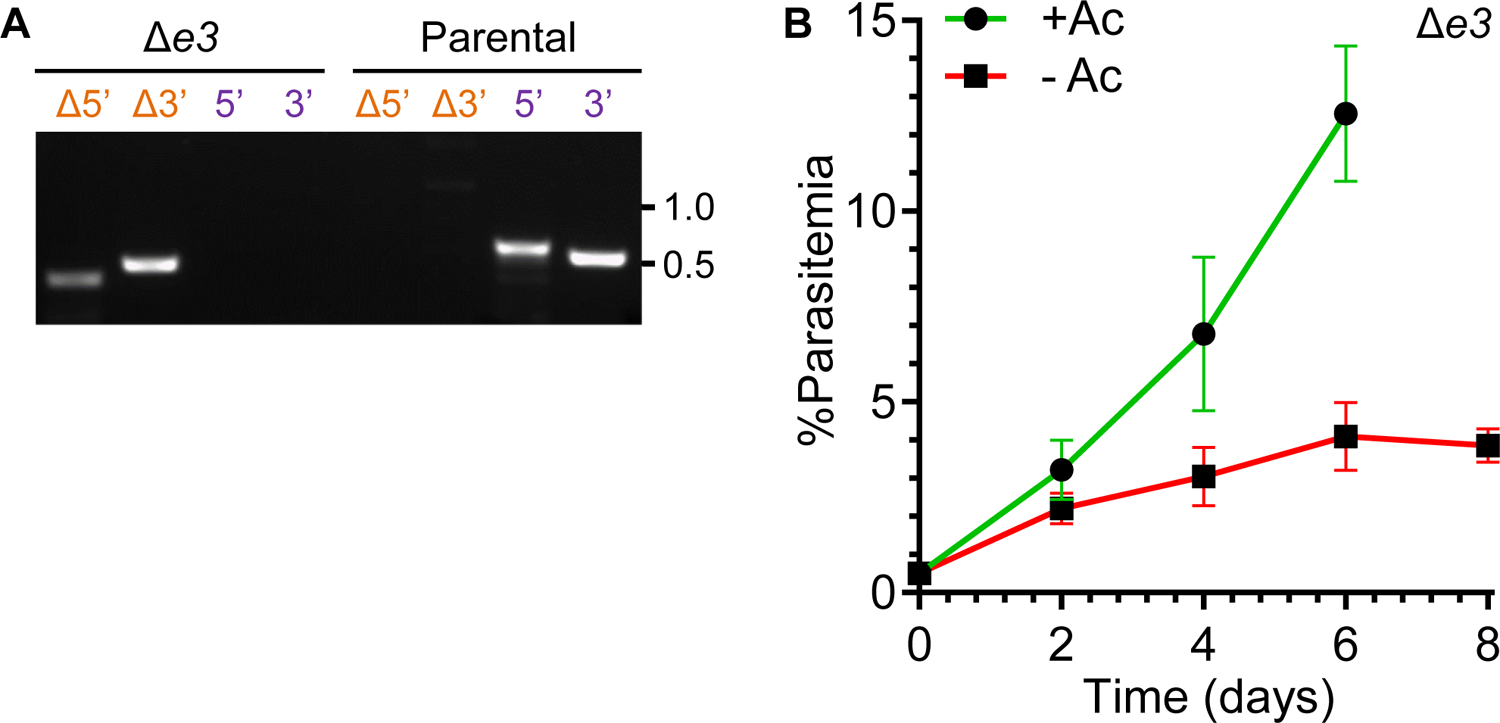
Uncropped gel images of clonal PCR analyses.

**Figure 2-figure supplement 1.**
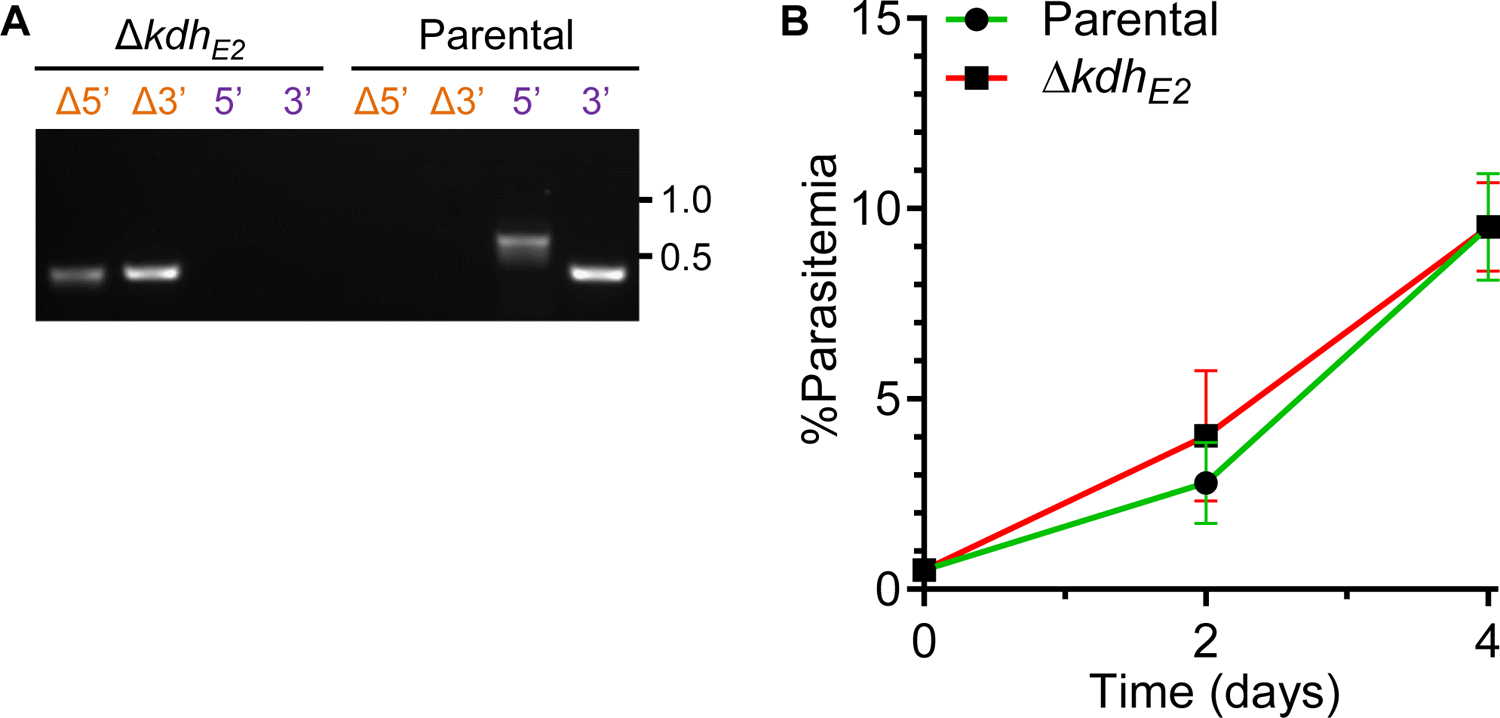
A. Genotyping PCR reactions confirming the deletion of the *e3* dehydrogenase gene. Based on the scheme shown in **Figure 1C**, PCR amplicons demonstrate integration at the Δ5’ and Δ3’ loci and lack of parental parasites (as indicated by the failure to amplify at the wild type 5’ and 3’ loci). The parental line was used as a control. The primer sequences and amplicon lengths are described in **supplementary file 1**. **B.** Growth curves comparing the growth of Δ*e3* parasites in the presence and absence of 5mM acetate (Ac). Error bars represent the standard deviation from two independent experiments, each conducted in quadruplicate.

**Figure 2-figure supplement 1-source data 1.** Uncropped gel images of clonal PCR analyses.

**Figure 3-figure supplement 1.**
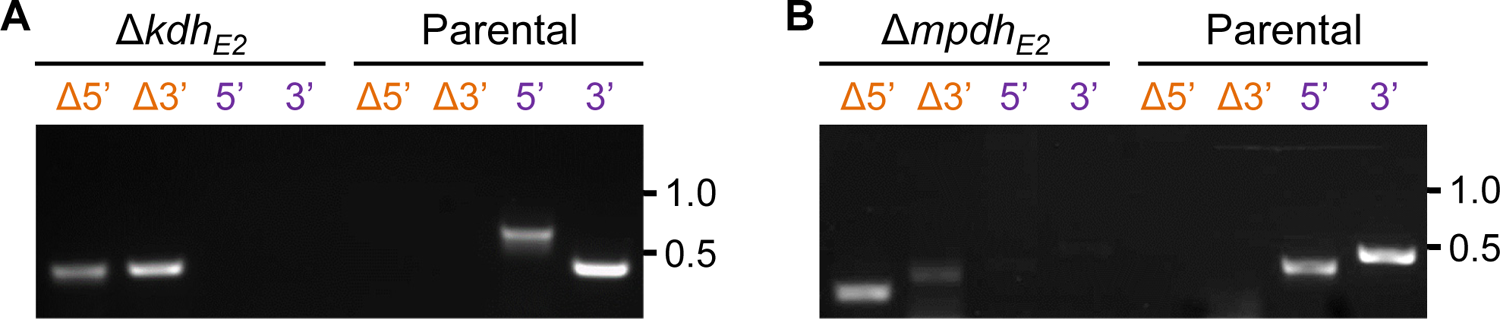
A. Genotyping PCR reactions confirming the deletion of the *kdh_E2_* gene. Based on the scheme shown in **Figure 1C**, PCR amplicons demonstrate integration at the Δ5’ and Δ3’ loci and lack of parental parasites (as indicated by the failure to amplify at the wild type 5’ and 3’ loci). The parental line was used as a control. The primer sequences and amplicon lengths are described in **supplementary file 1**. **B.** Growth curves of Δ*kdh_E2_* parasites (red) compared to the parental NF54^attB^ line (green). Error bars represent the standard deviation from two independent experiments, each conducted in quadruplicate.

**Figure 3-figure supplement 1-source data 1.** Uncropped gel images of clonal PCR analyses.

**Figure 3-figure supplement 2** A, B. Genotyping PCR reactions confirming the deletion of the *kdh_E2_* and *mpdh*_E2_ genes. Based on the scheme shown in **Figure 1C**, PCR amplicons demonstrate integration at the Δ5’ and Δ3’ loci and lack of parental parasites (as indicated by the failure to amplify at the wild type 5’ and 3’ loci). The parental line was used as a control. The primer sequences and amplicon lengths are described in **supplementary file 1**.

**Figure 3-figure supplement 2-source data 1.** Uncropped gel images of clonal PCR analyses.

**Figure 4-figure supplement 1.**
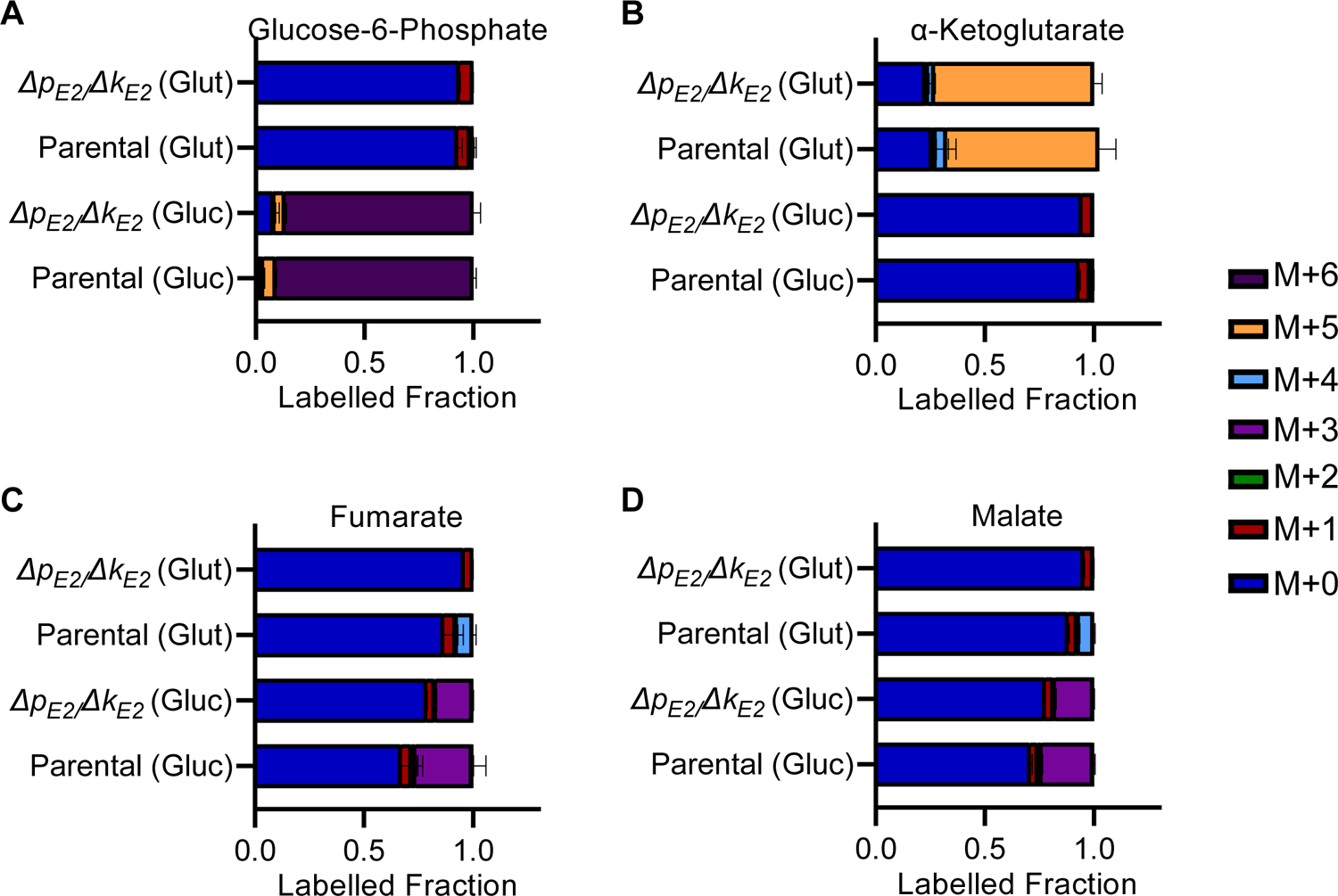
**A-D**. Fraction of isotopically labeled glucose 6-phosphate (**A**), ⍺-ketoglutarate (**B**), fumarate (**C**), or malate (**D**) when incubated with labeled glutamine (Glut) or glucose (Gluc) in parental or Δ*p_E2_/*Δ*k_E2_* parasites. For labeling experiments, color coding indicates the mass shift from the incorporation of heavy labeled carbon atoms in addition to the mass (M) of the parent compound. Labeling data are presented as the fraction of the total metabolite pool determined from N=3 experiments (parental) or N=2 experiments (Δ*p_E2_/*Δ*k_E2_*) with error bars representing the standard deviation (SD).

**Figure 5-figure supplement 1.**
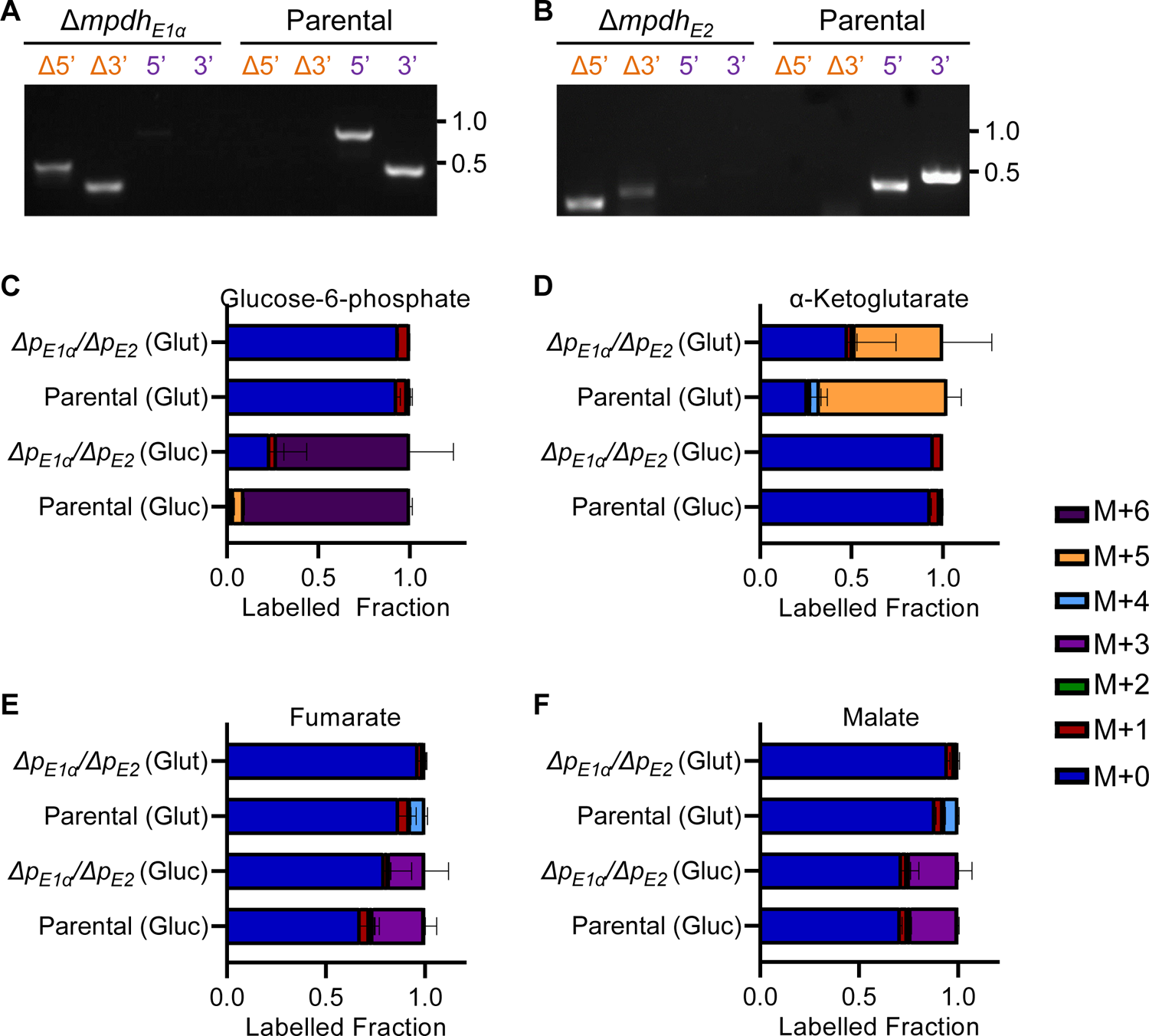
A, B. Genotyping PCR reactions confirming the deletion of the *mpdh_E1α_* and *mpdh*_E2_ genes. Based on the scheme shown in **Figure 1C**, PCR amplicons demonstrate integration at the Δ5’ and Δ3’ loci and lack of parental parasites (as indicated by the failure to amplify at the wild type 5’ and 3’ loci). The parental line was used as a control. The primer sequences and amplicon lengths are described in **supplementary file 1**. **C-F.** Fraction of isotopically labeled glucose-6-phosphate **(C)**, α-ketoglutarate **(D)**, fumarate **(E)**, or malate **(F)** when incubated with labeled glutamine (Glut) or glucose (Gluc) in parental or Δ*p*_E1α_*/*Δ*p*_E2_ parasites. For labeling experiments, color coding indicates the mass shift from the incorporation of heavy labeled carbon atoms in addition to the mass (M) of the parent compound. Labeling data are presented as the fraction of the total metabolite pool determined from N=3 experiments (parental) or N=2 experiments (Δ*p*_E1α_*/*Δ*p*_E2_) with error bars representing the standard deviation (SD).

**Figure 5-figure supplement 1-source data 1.** Uncropped gel images of clonal PCR analyses.

**Figure 6-figure supplement 1.**
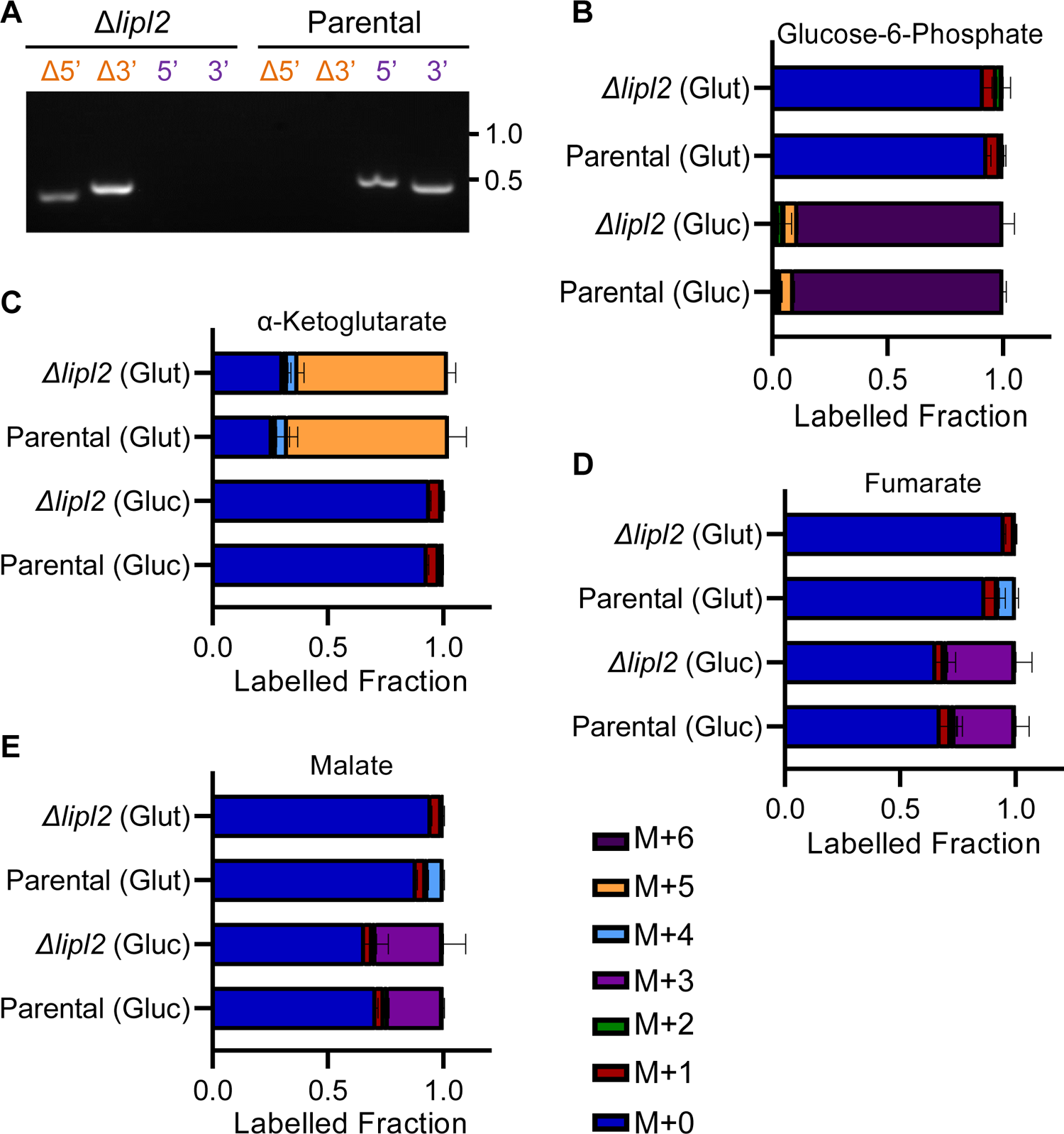
A. Genotyping PCR reactions confirming the deletion of the *lipl2* gene. Based on the scheme shown in **Figure 1C**, PCR amplicons demonstrate integration at the Δ5’ and Δ3’ loci and lack of parental parasites (as indicated by the failure to amplify at the wild type 5’ and 3’ loci). The parental line was used as a control. The primer sequences and amplicon lengths are described in **supplementary file 1**. **B-E.** Fraction of isotopically labeled glucose-6-phosphate **(B)**, α-ketoglutarate **(C)**, fumarate **(D)**, or malate **(E)** when incubated with labeled glutamine (Glut) or glucose (Gluc) in parental or *Δlipl2* parasites. For labeling experiments, color coding indicates the mass shift from the incorporation of heavy labeled carbon atoms in addition to the mass (M) of the parent compound. Labeling data are presented as the fraction of the total metabolite pool determined from N=3 experiments (parental) or N=2 experiments (*Δlipl2*) with error bars representing the standard deviation (SD).

**Figure 6-figure supplement 1-source data 1.** Uncropped gel images of clonal PCR analyses.

**Figure 7-source data 1.** Uncropped Western blot images demonstrating protein knockdown.

**Figure 7-figure supplement 1.**
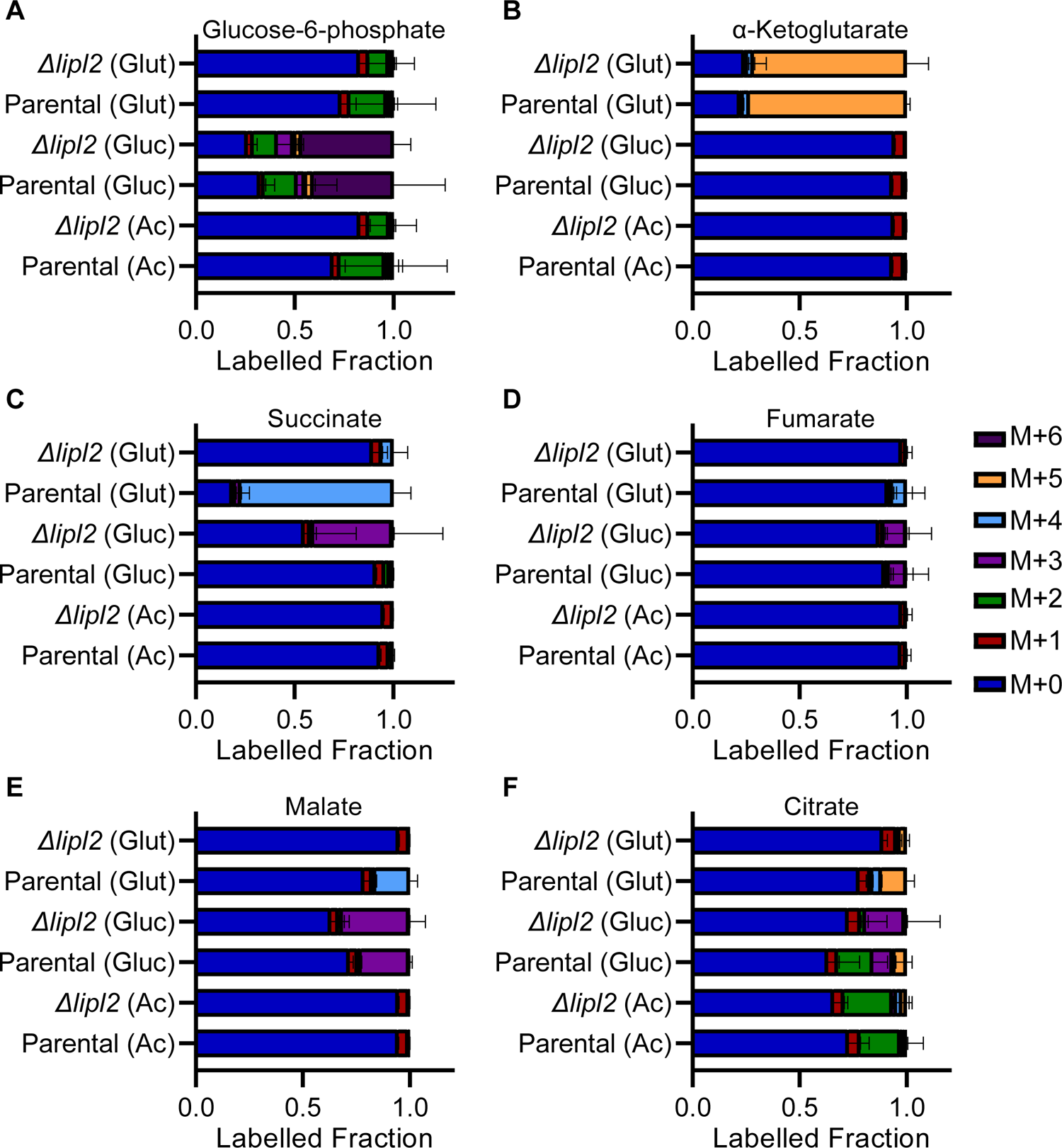
A-F. Fraction of isotopically labeled glucose-6-phosphate **(A)**, α-ketoglutarate **(B)**, succinate **(C)**, fumarate **(D)**, malate **(E)** or citrate **(F)** when incubated with labeled glutamine (Glut), glucose (Gluc) or acetate (Ac) in parental or Δ*lipl2* parasites. For labeling experiments, color coding indicates the mass shift from the incorporation of heavy labeled carbon atoms in addition to the mass (M) of the parent compound. Labeling data are presented as the fraction of the total metabolite pool determined from N=2 experiments with error bars representing the standard deviation (SD).

**Figure 7-figure supplement 2.**
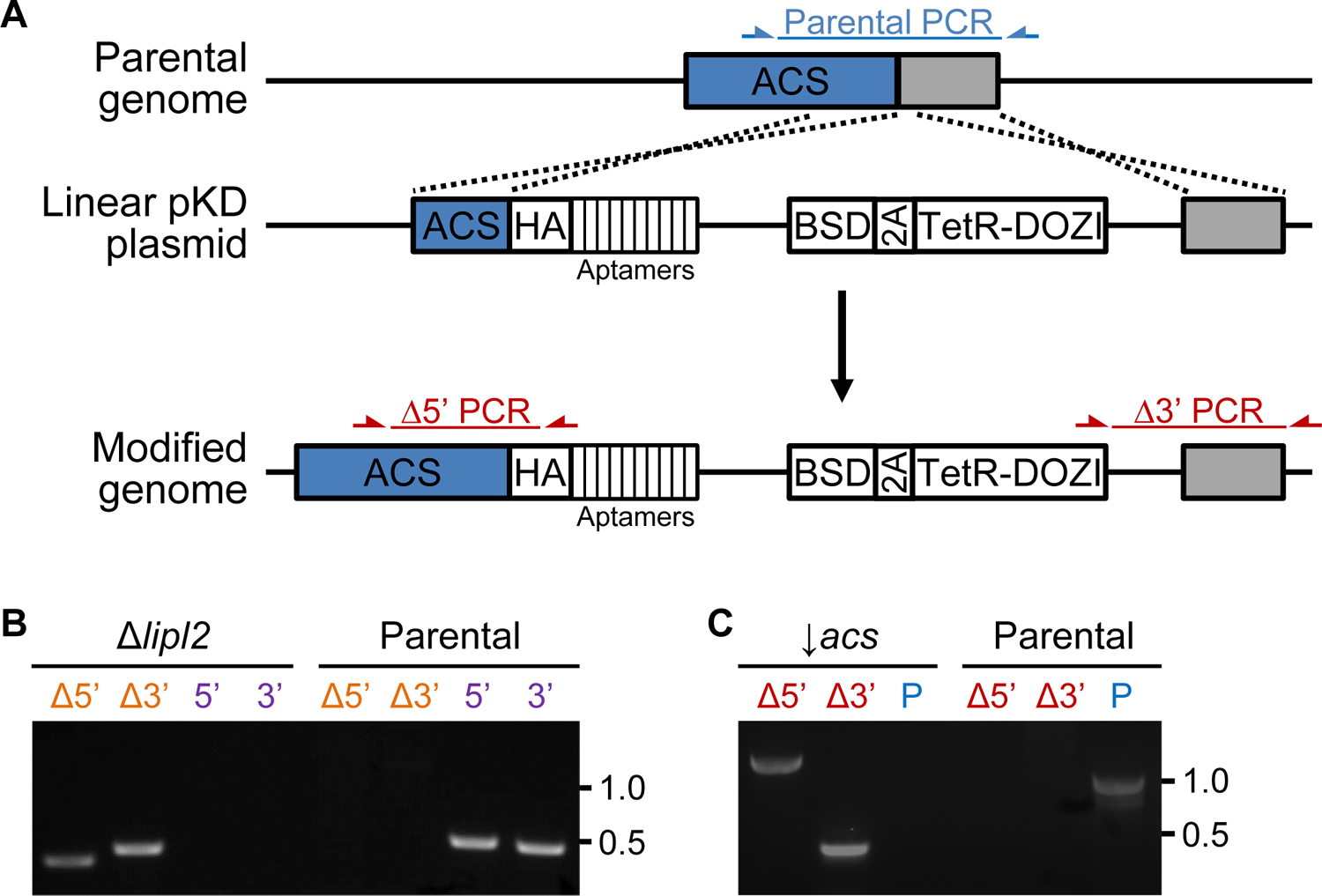
A. Schematic representation showing how the gene encoding acetyl-CoA synthetase (ACS) was modified using linearized pKD plasmid. Integration of the pKD plasmid appended a region encoding a c-terminal hemagglutinin tag (HA) and an array of ten aptamers. The modified genome also contained an expression cassette that produces tetracycline repressor (TetR) fused to the DOZI mRNA silencing protein (TetR-DOZI) and blasticidin-S deaminase (BSD). The positions of genotyping PCR products designed to identify the parental locus (blue) or the recombinant locus (red) are shown. The primer sequences and amplicon lengths are described in **supplementary file 1**. **B.** Genotyping PCR reactions confirming the deletion of the *lipl2* gene. Based on the scheme shown in **Figure 1C**, PCR amplicons demonstrate integration at the Δ5’ and Δ3’ loci and lack of parental parasites (as indicated by the failure to amplify at the wild type 5’ and 3’ loci). The parental line was used as a control. The primer sequences and amplicon lengths are described in **supplementary file 1**. **C.** Genotyping PCR reactions confirming the modification of the *acs* gene for knockdown experiments. Based on the scheme shown in **A**, PCR amplicons demonstrate integration at the Δ5’ and Δ3’ loci and lack of parental parasites (as indicated by the failure to amplify parental (P) locus). The parental line was used as a control. The primer sequences and amplicon lengths are described in **supplementary file 1**.

**Figure 7-figure supplement 2-source data 1.** Uncropped gel images of clonal PCR analyses.

**Figure 8-source data 1.** Uncropped Western blot images demonstrating protein knockdown.

**Figure 8-figure supplement 1.**
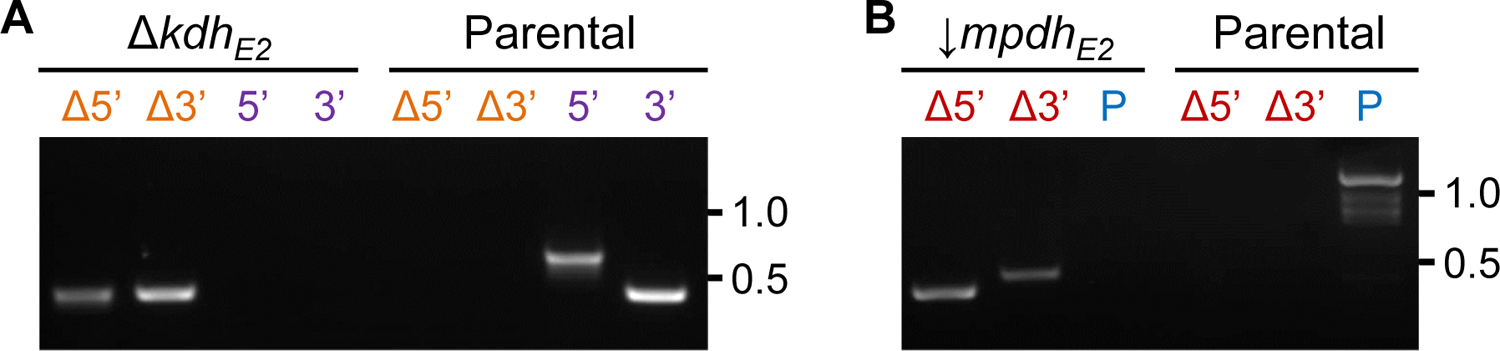
A. Genotyping PCR reactions confirming the deletion of the *kdh_E2_* gene. Based on the scheme shown in **Figure 1C**, PCR amplicons demonstrate integration at the Δ5’ and Δ3’ loci and lack of parental parasites (as indicated by the failure to amplify at the wild type 5’ and 3’ loci). The parental line was used as a control. The primer sequences and amplicon lengths are described in **supplementary file 1**. **B.** Genotyping PCR reactions confirming the modification of the *mpdh*_E2_ gene for knockdown experiments. Based on the scheme shown in **Figure 7-figure supplement 2A**, PCR amplicons demonstrate integration at the Δ5’ and Δ3’ loci and lack of parental parasites (as indicated by the failure to amplify parental (P) locus). The parental line was used as a control. The primer sequences and amplicon lengths are described in **supplementary file 1**.

**Figure 8-figure supplement 1-source data 1.** Uncropped gel images of clonal PCR analyses.

## Notes

### Competing Interest Statement

The authors have declared no competing interest.

## References

1. WHO. World Malaria Report. (2019).

2. Dhiman, S. Are malaria elimination efforts on right track? An analysis of gains achieved and challenges ahead. Infect Dis Poverty 8, 14, doi:10.1186/s40249-019-0524-x (2019).

3. Rts, S. C. T. P. Efficacy and safety of RTS,S/AS01 malaria vaccine with or without a booster dose in infants and children in Africa: final results of a phase 3, individually randomised, controlled trial. Lancet 386, 31–45, doi:10.1016/S0140-6736(15)60721-8 (2015).

4. Menard, D. & Dondorp, A. Antimalarial Drug Resistance: A Threat to Malaria Elimination. Cold Spring Harb Perspect Med 7, doi:10.1101/cshperspect.a025619 (2017).

5. Domingo, R. et al. Overcoming synthetic challenges in targeting coenzyme A biosynthesis with the antimicrobial natural product CJ-15,801. Medchemcomm 10, 2118–2125, doi:10.1039/c9md00312f (2019).

6. Schalkwijk, J. et al. Antimalarial pantothenamide metabolites target acetyl-coenzyme A biosynthesis in Plasmodium falciparum. Sci Transl Med 11, doi:10.1126/scitranslmed.aas9917 (2019).

7. Spry, C. et al. Structure-activity analysis of CJ-15,801 analogues that interact with Plasmodium falciparum pantothenate kinase and inhibit parasite proliferation. Eur J Med Chem 143, 1139–1147, doi:10.1016/j.ejmech.2017.08.050 (2018).

8. Fletcher, S. et al. Biological characterization of chemically diverse compounds targeting the Plasmodium falciparum coenzyme A synthesis pathway. Parasit Vectors 9, 589, doi:10.1186/s13071-016-1860-3 (2016).

9. Pietrocola, F., Galluzzi, L., Bravo-San Pedro, J. M., Madeo, F. & Kroemer, G. Acetyl coenzyme A: a central metabolite and second messenger. Cell Metab 21, 805–821, doi:10.1016/j.cmet.2015.05.014 (2015).

10. Shi, L. & Tu, B. P. Acetyl-CoA and the regulation of metabolism: mechanisms and consequences. Curr Opin Cell Biol 33, 125-131, doi:10.1016/j.ceb.2015.02.003 (2015).

11. Choudhary, C., Weinert, B. T., Nishida, Y., Verdin, E. & Mann, M. The growing landscape of lysine acetylation links metabolism and cell signalling. Nat Rev Mol Cell Biol 15, 536–550, doi:10.1038/nrm3841 (2014).

12. Trefely, S., Doan, M. T. & Snyder, N. W. Crosstalk between cellular metabolism and histone acetylation. Methods Enzymol 626, 1–21, doi:10.1016/bs.mie.2019.07.013 (2019).

13. Cobbold, S. A., Santos, J. M., Ochoa, A., Perlman, D. H. & Llinas, M. Proteome-wide analysis reveals widespread lysine acetylation of major protein complexes in the malaria parasite. Sci Rep 6, 19722, doi:10.1038/srep19722 (2016).

14. Cui, L. et al. PfGCN5-mediated histone H3 acetylation plays a key role in gene expression in Plasmodium falciparum. Eukaryot Cell 6, 1219–1227, doi:10.1128/EC.00062-07 (2007).

15. Rawat, M., Malhotra, R., Shintre, S., Pani, S. & Karmodiya, K. Role of PfGCN5 in nutrient sensing and transcriptional regulation in Plasmodium falciparum. J Biosci 45 (2020).

16. Chua, M. J. et al. Effect of clinically approved HDAC inhibitors on Plasmodium, Leishmania and Schistosoma parasite growth. Int J Parasitol Drugs Drug Resist 7, 42–50, doi:10.1016/j.ijpddr.2016.12.005 (2017).

17. Andrews, K. T., Tran, T. N. & Fairlie, D. P. Towards histone deacetylase inhibitors as new antimalarial drugs. Curr Pharm Des 18, 3467–3479 (2012).

18. Wheatley, N. C. et al. Antimalarial histone deacetylase inhibitors containing cinnamate or NSAID components. Bioorg Med Chem Lett 20, 7080–7084, doi:10.1016/j.bmcl.2010.09.096 (2010).

19. Andrews, K. T., Tran, T. N., Wheatley, N. C. & Fairlie, D. P. Targeting histone deacetylase inhibitors for anti-malarial therapy. Curr Top Med Chem 9, 292–308, doi:10.2174/156802609788085313 (2009).

20. Andrews, K. T. et al. Potent antimalarial activity of histone deacetylase inhibitor analogues. Antimicrob Agents Chemother 52, 1454–1461, doi:10.1128/AAC.00757-07 (2008).

21. Mackwitz, M. K. W. et al. Structure-Activity and Structure-Toxicity Relationships of Peptoid-Based Histone Deacetylase Inhibitors with Dual-Stage Antiplasmodial Activity. ChemMedChem 14, 912–926, doi:10.1002/cmdc.201800808 (2019).

22. Foth, B. J. et al. The malaria parasite Plasmodium falciparum has only one pyruvate dehydrogenase complex, which is located in the apicoplast. Mol Microbiol 55, 39–53, doi:10.1111/j.1365-2958.2004.04407.x (2005).

23. Pei, Y. et al. Plasmodium pyruvate dehydrogenase activity is only essential for the parasite’s progression from liver infection to blood infection. Mol Microbiol 75, 957–971, doi:10.1111/j.1365-2958.2009.07034.x (2010).

24. Prigge, S. T., He, X., Gerena, L., Waters, N. C. & Reynolds, K. A. The initiating steps of a type II fatty acid synthase in Plasmodium falciparum are catalyzed by pfACP, pfMCAT, and pfKASIII. Biochemistry 42, 1160–1169, doi:10.1021/bi026847k (2003).

25. Waters, N. C. et al. Functional characterization of the acyl carrier protein (PfACP) and beta-ketoacyl ACP synthase III (PfKASIII) from Plasmodium falciparum. Mol Biochem Parasitol 123, 85–94, doi:10.1016/s0166-6851(02)00140-8 (2002).

26. Vaughan, A. M. et al. Type II fatty acid synthesis is essential only for malaria parasite late liver stage development. Cell Microbiol 11, 506–520, doi:10.1111/j.1462-5822.2008.01270.x (2009).

27. Yu, M. et al. The fatty acid biosynthesis enzyme FabI plays a key role in the development of liver-stage malarial parasites. Cell Host Microbe 4, 567–578, doi:10.1016/j.chom.2008.11.001 (2008).

28. van Schaijk, B. C. et al. Type II fatty acid biosynthesis is essential for Plasmodium falciparum sporozoite development in the midgut of Anopheles mosquitoes. Eukaryot Cell 13, 550–559, doi:10.1128/EC.00264-13 (2014).

29. Cobbold, S. A. et al. Kinetic flux profiling elucidates two independent acetyl-CoA biosynthetic pathways in Plasmodium falciparum. J Biol Chem 288, 36338–36350, doi:10.1074/jbc.M113.503557 (2013).

30. Swift, R. P. et al. The NTP generating activity of pyruvate kinase II is critical for apicoplast maintenance in Plasmodium falciparum. Elife 9, doi:10.7554/eLife.50807 (2020).

31. Bryant, J. M. et al. Exploring the virulence gene interactome with CRISPR/dCas9 in the human malaria parasite. Mol Syst Biol 16, e9569, doi:10.15252/msb.20209569 (2020).

32. Prata, I. O. et al. Plasmodium falciparum Acetyl-CoA Synthetase is essential for parasite intraerythrocytic development and chromatin modification. ACS Infectious Diseases, 2021.2006.2013.448207, doi:10.1021/acsinfecdis.1c00414 (2021).

33. Summers, R. L. et al. Chemogenomics identifies acetyl-coenzyme A synthetase as a target for malaria treatment and prevention. Cell Chem Biol, doi:10.1016/j.chembiol.2021.07.010 (2021).

34. Oppenheim, R. D. et al. BCKDH: the missing link in apicomplexan mitochondrial metabolism is required for full virulence of Toxoplasma gondii and Plasmodium berghei. PLoS Pathog 10, e1004263, doi:10.1371/journal.ppat.1004263 (2014).

35. Spalding, M. D. & Prigge, S. T. Lipoic acid metabolism in microbial pathogens. Microbiol Mol Biol Rev 74, 200–228, doi:10.1128/MMBR.00008-10 (2010).

36. Wrenger, C. & Muller, S. The human malaria parasite Plasmodium falciparum has distinct organelle-specific lipoylation pathways. Mol Microbiol 53, 103–113, doi:10.1111/j.1365-2958.2004.04112.x (2004).

37. Falkard, B. et al. A key role for lipoic acid synthesis during Plasmodium liver stage development. Cell Microbiol 15, 1585–1604, doi:10.1111/cmi.12137 (2013).

38. Gunther, S. et al. Apicoplast lipoic acid protein ligase B is not essential for Plasmodium falciparum. PLoS Pathog 3, e189, doi:10.1371/journal.ppat.0030189 (2007).

39. Biddau, M. et al. Plasmodium falciparum LipB mutants display altered redox and carbon metabolism in asexual stages and cannot complete sporogony in Anopheles mosquitoes. Int J Parasitol 51, 441–453, doi:10.1016/j.ijpara.2020.10.011 (2021).

40. Allary, M., Lu, J. Z., Zhu, L. & Prigge, S. T. Scavenging of the cofactor lipoate is essential for the survival of the malaria parasite Plasmodium falciparum. Mol Microbiol 63, 1331–1344, doi:10.1111/j.1365-2958.2007.05592.x (2007).

41. Afanador, G. A. et al. Redox-dependent lipoylation of mitochondrial proteins in Plasmodium falciparum. Mol Microbiol 94, 156–171, doi:10.1111/mmi.12753 (2014).

42. Jhun, H., Walters, M. S. & Prigge, S. T. Using Lipoamidase as a Novel Probe To Interrogate the Importance of Lipoylation in Plasmodium falciparum. mBio 9, doi:10.1128/mBio.01872-18 (2018).

43. McMillan, P. J., Stimmler, L. M., Foth, B. J., McFadden, G. I. & Muller, S. The human malaria parasite Plasmodium falciparum possesses two distinct dihydrolipoamide dehydrogenases. Mol Microbiol 55, 27–38, doi:10.1111/j.1365-2958.2004.04398.x (2005).

44. Bushell, E. et al. Functional Profiling of a Plasmodium Genome Reveals an Abundance of Essential Genes. Cell 170, 260–272 e268, doi:10.1016/j.cell.2017.06.030 (2017).

45. Zhang, M. et al. Uncovering the essential genes of the human malaria parasite Plasmodium falciparum by saturation mutagenesis. Science 360, doi:10.1126/science.aap7847 (2018).

46. Ke, H. et al. Genetic investigation of tricarboxylic acid metabolism during the Plasmodium falciparum life cycle. Cell Rep 11, 164–174, doi:10.1016/j.celrep.2015.03.011 (2015).

47. Salcedo, E., Sims, P. F. & Hyde, J. E. A glycine-cleavage complex as part of the folate one-carbon metabolism of Plasmodium falciparum. Trends Parasitol 21, 406–411, doi:10.1016/j.pt.2005.07.001 (2005).

48. Spalding, M. D., Allary, M., Gallagher, J. R. & Prigge, S. T. Validation of a modified method for Bxb1 mycobacteriophage integrase-mediated recombination in Plasmodium falciparum by localization of the H-protein of the glycine cleavage complex to the mitochondrion. Mol Biochem Parasitol 172, 156–160, doi:10.1016/j.molbiopara.2010.04.005 (2010).

49. Chan, X. W. et al. Chemical and genetic validation of thiamine utilization as an antimalarial drug target. Nat Commun 4, 2060, doi:10.1038/ncomms3060 (2013).

50. Tian, J. et al. Mycobacterium tuberculosis appears to lack alpha-ketoglutarate dehydrogenase and encodes pyruvate dehydrogenase in widely separated genes. Mol Microbiol 57, 859–868, doi:10.1111/j.1365-2958.2005.04741.x (2005).

51. Afanador, G. A. et al. A novel lipoate attachment enzyme is shared by Plasmodium and Chlamydia species. Mol Microbiol 106, 439–451, doi:10.1111/mmi.13776 (2017).

52. Ganesan, S. M., Falla, A., Goldfless, S. J., Nasamu, A. S. & Niles, J. C. Synthetic RNA-protein modules integrated with native translation mechanisms to control gene expression in malaria parasites. Nat Commun 7, 10727, doi:10.1038/ncomms10727 (2016).

53. Rajaram, K., Liu, H. B. & Prigge, S. T. Redesigned TetR-Aptamer System To Control Gene Expression in Plasmodium falciparum. mSphere 5, doi:10.1128/mSphere.00457-20 (2020).

54. Kocherginsky, N. Acidic lipids, H(+)-ATPases, and mechanism of oxidative phosphorylation. Physico-chemical ideas 30 years after P. Mitchell’s Nobel Prize award. Prog Biophys Mol Biol 99, 20-41, doi:10.1016/j.pbiomolbio.2008.10.013 (2009).

55. Sturm, A., Mollard, V., Cozijnsen, A., Goodman, C. D. & McFadden, G. I. Mitochondrial ATP synthase is dispensable in blood-stage Plasmodium berghei rodent malaria but essential in the mosquito phase. Proc Natl Acad Sci U S A 112, 10216–10223, doi:10.1073/pnas.1423959112 (2015).

56. Painter, H. J., Morrisey, J. M., Mather, M. W. & Vaidya, A. B. Specific role of mitochondrial electron transport in blood-stage Plasmodium falciparum. Nature 446, 88–91, doi:10.1038/nature05572 (2007).

57. Palmer, M. J. et al. Potent Antimalarials with Development Potential Identified by Structure-Guided Computational Optimization of a Pyrrole-Based Dihydroorotate Dehydrogenase Inhibitor Series. J Med Chem 64, 6085–6136, doi:10.1021/acs.jmedchem.1c00173 (2021).

58. Kokkonda, S. et al. Lead Optimization of a Pyrrole-Based Dihydroorotate Dehydrogenase Inhibitor Series for the Treatment of Malaria. J Med Chem 63, 4929–4956, doi:10.1021/acs.jmedchem.0c00311 (2020).

59. Vaidya, A. B. & Mather, M. W. Mitochondrial evolution and functions in malaria parasites. Annu Rev Microbiol 63, 249–267, doi:10.1146/annurev.micro.091208.073424 (2009).

60. Mathur, V., Wakeman, K. C. & Keeling, P. J. Parallel functional reduction in the mitochondria of apicomplexan parasites. Curr Biol 31, 2920–2928 e2924, doi:10.1016/j.cub.2021.04.028 (2021).

61. Dellibovi-Ragheb, T. A., Gisselberg, J. E. & Prigge, S. T. Parasites FeS up: iron-sulfur cluster biogenesis in eukaryotic pathogens. PLoS Pathog 9, e1003227, doi:10.1371/journal.ppat.1003227 (2013).

62. Kloehn, J. et al. Multi-omics analysis delineates the distinct functions of sub-cellular acetyl-CoA pools in Toxoplasma gondii. BMC Biol 18, 67, doi:10.1186/s12915-020-00791-7 (2020).

63. Tymoshenko, S. et al. Metabolic Needs and Capabilities of Toxoplasma gondii through Combined Computational and Experimental Analysis. PLoS Comput Biol 11, e1004261, doi:10.1371/journal.pcbi.1004261 (2015).

64. Dolce, V., Cappello, A. R. & Capobianco, L. Mitochondrial tricarboxylate and dicarboxylate-tricarboxylate carriers: from animals to plants. IUBMB Life 66, 462–471, doi:10.1002/iub.1290 (2014).

65. van Rossum, H. M., Kozak, B. U., Pronk, J. T. & van Maris, A. J. A. Engineering cytosolic acetyl-coenzyme A supply in Saccharomyces cerevisiae: Pathway stoichiometry, free-energy conservation and redox-cofactor balancing. Metab Eng 36, 99–115, doi:10.1016/j.ymben.2016.03.006 (2016).

66. Wei, X., Schultz, K., Bazilevsky, G. A., Vogt, A. & Marmorstein, R. Molecular basis for acetyl-CoA production by ATP-citrate lyase. Nat Struct Mol Biol 27, 33–41, doi:10.1038/s41594-019-0351-6 (2020).

67. Nozawa, A. et al. Characterization of mitochondrial carrier proteins of malaria parasite Plasmodium falciparum based on in vitro translation and reconstitution. Parasitol Int 79, 102160, doi:10.1016/j.parint.2020.102160 (2020).

68. Nozawa, A., Fujimoto, R., Matsuoka, H., Tsuboi, T. & Tozawa, Y. Cell-free synthesis, reconstitution, and characterization of a mitochondrial dicarboxylate-tricarboxylate carrier of Plasmodium falciparum. Biochem Biophys Res Commun 414, 612–617, doi:10.1016/j.bbrc.2011.09.130 (2011).

69. Madiraju, P., Pande, S. V., Prentki, M. & Madiraju, S. R. Mitochondrial acetylcarnitine provides acetyl groups for nuclear histone acetylation. Epigenetics 4, 399–403, doi:10.4161/epi.4.6.9767 (2009).

70. Bieber, L. L. Carnitine. Annu Rev Biochem 57, 261–283, doi:10.1146/annurev.bi.57.070188.001401 (1988).

71. Fleck, C. B. & Brock, M. Re-characterisation of Saccharomyces cerevisiae Ach1p: fungal CoA-transferases are involved in acetic acid detoxification. Fungal Genet Biol 46, 473–485, doi:10.1016/j.fgb.2009.03.004 (2009).

72. Flikweert, M. T., de Swaaf, M., van Dijken, J. P. & Pronk, J. T. Growth requirements of pyruvate-decarboxylase-negative Saccharomyces cerevisiae. FEMS Microbiol Lett 174, 73–79, doi:10.1111/j.1574-6968.1999.tb13551.x (1999).

73. Van den Berg, M. A. & Steensma, H. Y. ACS2, a Saccharomyces cerevisiae gene encoding acetyl-coenzyme A synthetase, essential for growth on glucose. Eur J Biochem 231, 704–713, doi:10.1111/j.1432-1033.1995.tb20751.x (1995).

74. Lim, M. Y. et al. UDP-galactose and acetyl-CoA transporters as Plasmodium multidrug resistance genes. Nat Microbiol 1, 16166, doi:10.1038/nmicrobiol.2016.166 (2016).

75. Boateng, R. A., Tastan Bishop, O. & Musyoka, T. M. Characterisation of plasmodial transketolases and identification of potential inhibitors: an in silico study. Malar J 19, 442, doi:10.1186/s12936-020-03512-1 (2020).

76. Tollinger, C. D., Vreman, H. J. & Weiner, M. W. Measurement of acetate in human blood by gas chromatography: effects of sample preparation, feeding, and various diseases. Clin Chem 25, 1787–1790 (1979).

77. Ghorbal, M. et al. Genome editing in the human malaria parasite Plasmodium falciparum using the CRISPR-Cas9 system. Nat Biotechnol 32, 819–821, doi:10.1038/nbt.2925 (2014).

78. Swift, R. P. et al. A mevalonate bypass system facilitates elucidation of plastid biology in malaria parasites. PLoS Pathog 16, e1008316, doi:10.1371/journal.ppat.1008316 (2020).

79. Swift, R. P., Rajaram, K., Liu, H. B. & Prigge, S. T. Dephospho-CoA kinase, a nuclear-encoded apicoplast protein, remains active and essential after Plasmodium falciparum apicoplast disruption. EMBO J 40, e107247, doi:10.15252/embj.2020107247 (2021).

80. Keller, A., Nesvizhskii, A. I., Kolker, E. & Aebersold, R. Empirical statistical model to estimate the accuracy of peptide identifications made by MS/MS and database search. Anal Chem 74, 5383–5392, doi:10.1021/ac025747h (2002).

81. Nesvizhskii, A. I., Keller, A., Kolker, E. & Aebersold, R. A statistical model for identifying proteins by tandem mass spectrometry. Anal Chem 75, 4646–4658, doi:10.1021/ac0341261 (2003).

82. Agrawal, S. et al. El-MAVEN: A Fast, Robust, and User-Friendly Mass Spectrometry Data Processing Engine for Metabolomics. Methods Mol Biol 1978, 301–321, doi:10.1007/978-1-4939-9236-2_19 (2019).

